# CAR T cell targeting of inflammatory myeloid progenitors in the bone marrow remodels border-associated macrophages and reverses cognitive aging

**DOI:** 10.64898/2026.09.27.754777

**Authors:** Alexander S. Harris, James A. Rouse, Guangran Guo, Inés Fernandez-Maestre, Sujay Pal, Brian Oh, Shuxuan Li, Wen Yi See, Arianna Anderson, Joseph Gewolb, Matthew Soethout, Jill Habel, Rad Utama, Christopher R. Vakoc, Corina Amor

## Abstract

Aging is associated with progressive neuroinflammation and cognitive decline, yet the cellular sources that sustain this process and whether they can be targeted peripherally remain unclear. Here, we identify inflammatory border-associated macrophages (BAMs) as key drivers of neuroinflammation in aging and show that their therapeutic and prophylactic elimination through intrathecal CAR T cell therapy restores cognitive performance in mouse models of aging and Alzheimer’s disease. Furthermore, targeting of aged inflammatory bone marrow myeloid progenitors through either intravenous CAR T cells, which do not infiltrate the brain, or through transplantation of CAR T-treated progenitors is sufficient to reduce neuroinflammation and cognitive impairment. These findings reveal that bone marrow myeloid progenitors harbor a heritable inflammatory transcriptional state that is transmitted to their BAM progeny, driving neuroinflammation and cognitive deterioration, and conserved in human aging. Critically, this proinflammatory state is marked by the upregulation of surface proteins, enabling precise peripheral CAR T cell targeting of these progenitors for long-lasting therapeutic effects in cognitive aging.

## Introduction

Aging is the primary risk factor for the development of cognitive decline and neurodegenerative conditions such as Alzheimer’s disease (AD)^1^. Current therapeutic approaches have focused on targeting downstream elements of the pathology such as amyloid-β plaques or tau neurofibrillary tangles through monoclonal antibodies^2,3^. These have limited long-term efficacy and require continuous repeated dosing, highlighting the need for novel therapeutic strategies.

Neuroinflammation is a hallmark of both aging and neurodegeneration, and has been implicated as a driver of cognitive decline^4^. Therapeutic strategies targeting neuroinflammation currently in clinical development include broad chemical approaches to modulate inflammation such as tyrosine kinase inhibitors and cytokine-targeting agents, as well as microglial modulators^5^. These approaches focus on broadly suppressing neuroinflammation or targeting brain resident immune cells. However, whether there are upstream peripheral cellular drivers that sustain neuroinflammation in aging and whether they can be precisely targeted remains unclear.

In homeostatic conditions, the immune landscape in mouse and human brains is primarily composed of microglia and border associated macrophages (BAMs) also referred to as CNS-associated macrophages (CAMs)^6–8^. Located at the brain’s border tissues (such as meninges, choroid plexus, and perivascular spaces), BAMs play key roles in immune defense, glymphatic clearance and modulation of extracellular matrix composition^6,7^. However, whether BAMs are causally responsible for neuroinflammation and cognitive decline in aging and tauopathy remains unclear. While initially derived from the yolk sac, emerging evidence suggests that disruption of brain homeostasis can lead to the partial replenishment of BAMs by bone marrow-derived monocytes, with these BAMs being transcriptionally distinct from yolk sac derived counterparts^9–12^. Whether peripheral mechanisms contribute to dysfunctional BAM phenotypes in aging, and whether these can be therapeutically targeted through peripheral approaches to alleviate neuroinflammation and cognitive decline in aging remain unknown.

Initially developed for refractory hematological malignancies, chimeric antigen receptor (CAR) T cells are showing remarkable potential in non-oncological indications including age-related dysfunction in peripheral tissues, where single administration of CAR T cells targeting uPAR (a surface marker of dysfunctional proinflammatory cells) leads to long-term improvements^13,14^. Recent studies have begun to explore the applicability of CARs for neurodegeneration by employing CD4 CAR T or CAR astrocytes targeting amyloid-β plaques^15,16^. However, beyond targeting pathological protein aggregates, the development of CARs for neurodegeneration has been hindered by the lack of suitable cell targets. Identifying upstream cellular drivers of neuroinflammation could thus open the door to the first cell-targeting CAR T approaches for neurodegeneration.

Here, using CAR T cells targeting the inflammatory cell marker uPAR as biological probe in mouse models of aging and tauopathy, we identified a subset of inflammatory BAMs as key drivers of neuroinflammation and cognitive impairment. These BAMs arose from bone marrow inflammatory myeloid progenitors that upregulated uPAR surface expression and heritably transmitted their proinflammatory state to their progeny. Peripheral targeting of these proinflammatory progenitors through single dose intravenous CAR T cells suppressed proinflammatory BAMs, resulting in decreased neuroinflammation and improved cognitive function. Taken together, these findings reveal a peripheral cellular target for CAR T therapy that reverses neuroinflammation and cognitive decline in aging and neurodegeneration.

## Results

### Intrathecal uPAR CAR T cells improve and prevent cognitive impairment in aging and tauopathy

To identify the optimal route of administration of uPAR CAR T cells to reach the brains of old animals, we compared cell infiltration when administered intravenously vs intrathecally **(Figure 1A)**. A single intrathecal, but not intravenous, administration of 1×10^6^ m.uPAR-m.28z or untransduced T (UT) cells generated from CD45.1 mice resulted in significant uPAR CAR T numbers in the brains of aged CD45.2 mice as determined by flow cytometry 20 days post infusion **(Figure 1B)**. The CAR T cells displayed an effector phenotype and their accumulation in the brain was well tolerated and did not result in clinical signs of toxicity such as weight loss or temperature changes **(Figure S1A-E)**. Intrathecally administered CAR T cells presented long term persistence, being still detected in the brains of animals 16 months after one infusion in their youth at age 3 months **(Figure 1C and S1F-G)**.

**Figure 1.**
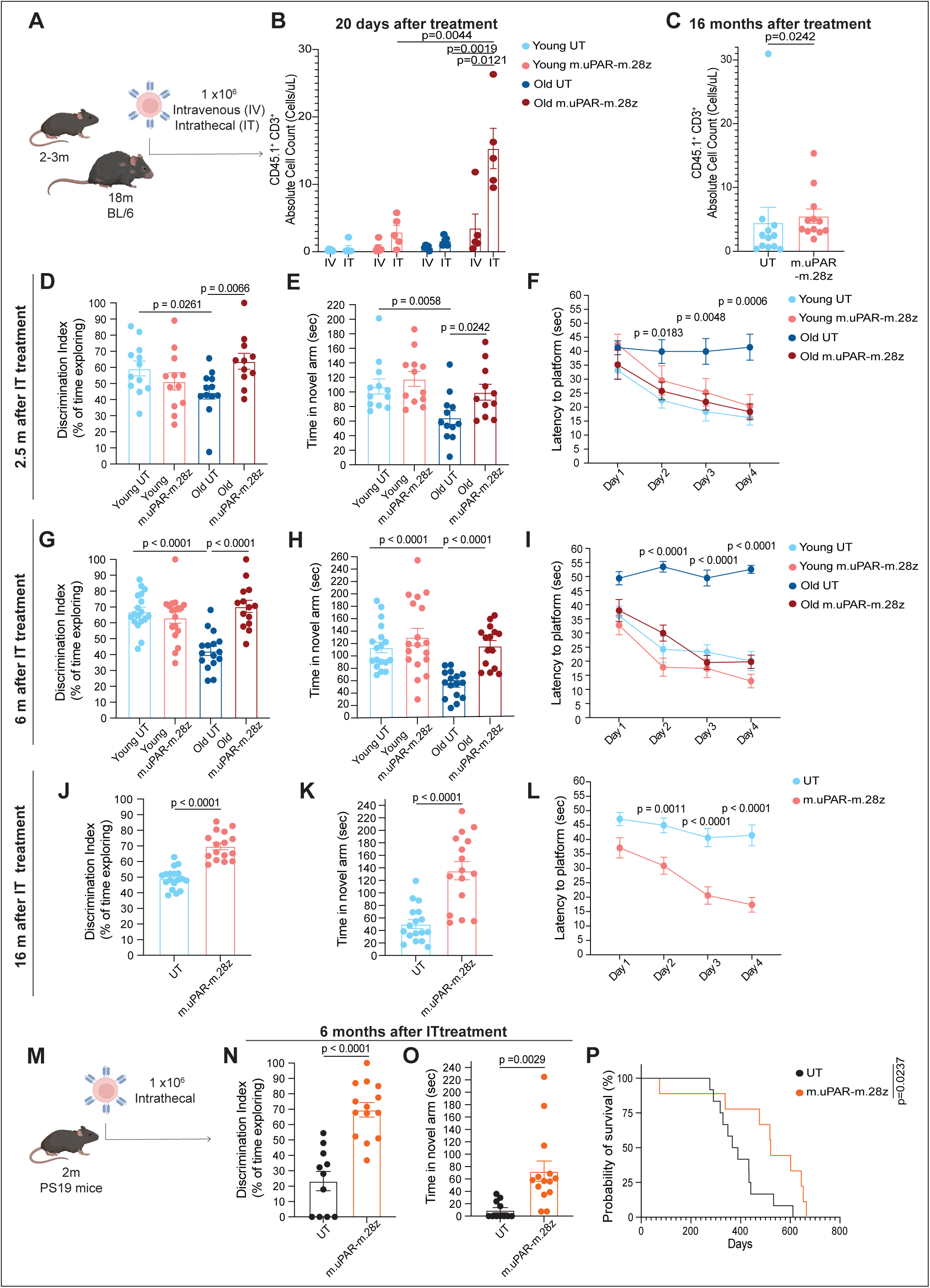
Intrathecal uPAR CAR T cells improve and prevent cognitive decline in aging and Tauopathy. (A) Schematic of experimental design for (B-L): Young (2-3 months) and old (18 months) mice were infused intravenously (IV) or intrathecally (IT) with either 1×10^6^ untransduced T cells (UT) or m.uPAR-m.28z CAR T cells from CD45.1 mice. (B) Absolute counts of CD45.1 and CD3 double positive cells in the brain as determined by flow cytometry 20 days after cell infusion. (n=5 mice per group). (C) Absolute counts of CD45.1 and CD3 double positive cells in the brain as determined by flow cytometry 16 months after intrathecal cell infusion. (n=12 mice per group). (D) Discrimination Index in the novel object recognition task 2.5m after intrathecal cell infusion. (UT young, m.uPAR-m.28z young and UT old: n=12 mice per group; m.uPAR-m.28z old: n=11 mice). (E) Time spent in novel arm in the Y-maze spatial memory task 2.5m after intrathecal cell infusion (UT young, m.uPAR-m.28z young and UT old: n=12 mice per group; m.uPAR-m.28z old: n=11 mice). (F) Latency to the platform in the Morris water maze test 2.5m after intrathecal cell infusion. (UT young, m.uPAR-m.28z young and UT old: n=12 mice per group; m.uPAR-m.28z old: n=10 mice). (G) Discrimination Index in the novel object recognition task 6m after intrathecal cell infusion. (UT young, m.uPAR-m.28z young: n=18 mice per group, UT old: n=16 mice; m.uPAR-m.28z old: n=14 mice). (H) Time spent in novel arm in the Y-maze spatial memory task 6m after intrathecal cell infusion. (UT young, m.uPAR-m.28z young: n=18 mice per group, UT old: n=16 mice; m.uPAR-m.28z old: n=15 mice). (I) Latency to the platform in the Morris water maze test 6m after intrathecal cell infusion. (UT young, m.uPAR-m.28z young: n= 12 mice per group, UT old: n=10 mice; m.uPAR-m.28z old: n=11 mice). (J) Discrimination Index in the novel object recognition task 16m after intrathecal cell infusion. (UT n=18 mice, m.uPAR-m.28z n=16 mice). (K) Time spent in novel arm in the Y-maze spatial memory task 16m after intrathecal cell infusion. (UT n=17 mice, m.uPAR-m.28z n=16 mice). (L) Latency to the platform in the Morris water maze test 16m after intrathecal cell infusion. (UT n=17 mice, m.uPAR-m.28z n=16 mice). (M) Schematic of experimental design for (N-P): PS19 mice (2-2.5 month) were infused intrathecally (IT) with 1×10^6^ untransduced T cells (UT) or m.uPAR-m.28z CAR T cells. (N) Discrimination Index in the novel object recognition task 6m after intrathecal cell infusion. (UT n=11 mice, m.uPAR-m.28z n=14 mice). (O) Time spent in novel arm in the Y-maze spatial memory task 6m after intrathecal cell infusion. (UT n=11 mice, m.uPAR-m.28z n=14 mice). (P) Kaplan-Meier survival curve of PS19 mice infused with intrathecally (IT) with 1×10^6^ untransduced T cells (UT) or m.uPAR-m.28z CAR T cells. (UT n=12 mice, m.uPAR-m.28z n=9 mice). Data are the mean ± s.e.m. (B)-(L) and (N)-(O). Statistical analysis was performed using two-tailed unpaired Student’s t-test (B) and (D-L) and (N-O); Mann-Whitney test (C) and Log-rank (Mantel-Cox) test (P). Data (B), (D-E) and (P) represent one independent experiment or (C) and (F-L) two independent experiments or (N-O) three independent experiments.

To examine the effect of intrathecal uPAR CAR T cells on age-related cognitive impairment, we performed: novel object tests to probe recognition memory, Y-maze spontaneous alternation to test spatial working memory, and Morris water maze to assess spatial learning and memory. Across these three different cognitive tests, old animals treated with intrathecal uPAR CAR T cells significantly outperformed old control treated mice **(Figure 1D-I and S1H-M)**. This cognitive rescue of age-related cognitive impairment was observed 2.5 months after a single intrathecal administration and was sustained long-term, even 6 months after treatment **(Figure 1D-I and S1H-M)**. Furthermore, young animals that had been prophylactically treated at 2 months of age, presented retained learning and memory function upon reaching 18 months of age (16 months after a single cell infusion) **(Figure 1J-L and S1N-P)**.

Beyond aging, we examined whether uPAR CAR T cells could potentially improve cognitive function in the PS19 transgenic mouse model, which expresses human MAPT^301S^ and develops robust neuroinflammation, cognitive impairment and premature mortality providing a stringent setting for testing the impact of disease-modifying interventions^17^ **(Figure 1M)**. A single intrathecal administration of 1×10^6^ uPAR CAR T cells at 2.5-3 months of age attenuated cognitive impairment **(Figure 1N-O and S1Q-R)** and resulted in a 37.8% increase in median survival **(Figure 1P)**.

Taken together, these results suggest that intrathecal administration of uPAR CAR T cells is well tolerated and can both rescue and attenuate age and tau-driven cognitive impairment.

### Intrathecal CAR T cells target inflammatory uPAR^+^ BAMs

To elucidate the identity of the cells being targeted by intrathecal uPAR CAR T cells in aged brains, we analyzed the surface expression of uPAR in this tissue using flow cytometry and immunofluorescence. uPAR levels significantly increased with age in mouse brains, particularly in immune populations **(Figure S2A-B)**. To identify their nature, we isolated through fluorescence-activated cell sorting (FACS) surface uPAR^+^ and uPAR^−^ cells from aged (18 month old) mouse brains and performed single-cell RNA sequencing (scRNAseq) profiling 8,941 uPAR^+^ and 20,177 uPAR^−^ individual cells **(Figure 2A)**. Using unsupervised clustering and the Allen Brain MapMyCells annovation tool^18^, we assigned 14 different cell types and visualized them with Uniform Manifold Approximation and Projection (UMAP) **(Figure S2C)**. Analysis of these different populations revealed that the majority of uPAR^+^ cells in aged brains were border associated macrophages (BAMs), which we further confirmed at the histological level observing a significant increase in the number of uPAR^+^ CD206^+^ cells in the brains of 18 months old mice compared with 3 months old mice **(Figure 2B and S2D-E)**. To understand their nature, we performed pathway analysis on the differentially expressed genes between uPAR^+^ and uPAR^−^ cells and found that uPAR^+^ cells in aged brains, and especially uPAR^+^ BAMs, were significantly enriched in terms related to the inflammatory response and neuroinflammation **(Figure 2C-D)**.

**Figure 2.**
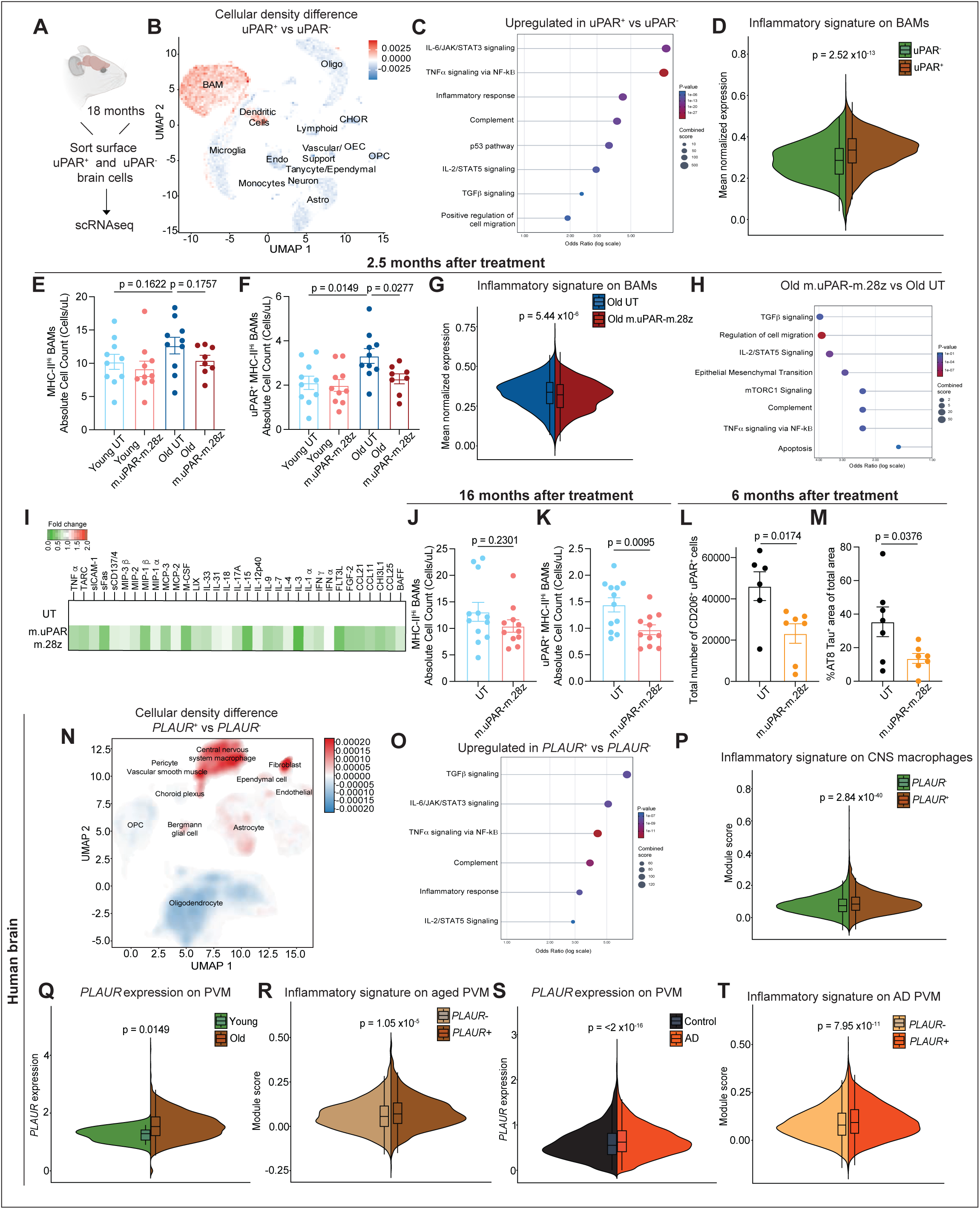
Intrathecal uPAR CAR T cells target inflammatory BAMs. (A) Experimental scheme for B-D: uPAR^+^ and uPAR^−^ cells from the brain of old (18 months old) mice were FACS sorted and subjected to scRNA-seq (n=12 mice pooled into two replicates). (B) UMAP-based binned density-difference map of retained aged-brain cells; density was calculated by binning the UMAP and normalizing cell counts within each uPAR group. Color indicates the relative local enrichment of uPAR^+^ versus uPAR^−^ cells. Labels denote 14 broad cell types assigned with MapMyCells using the 10x Whole Mouse Brain taxonomy^18,39^ (n=12 mice pooled into two replicates). (C) Enrichr over-representation analysis^43^ of direction-separated genes differentially expressed between uPAR^+^ and uPAR^−^ cells. Odds ratio is shown on the x axis, nominal enrichment P value by color and Enrichr combined score by point size (n=12 mice pooled into two replicates). (D) Split-violin plot of the mean LogNormalized expression of the available genes from the myeloid inflammatory signature^42^ in uPAR^−^ and uPAR^+^ BAMs. Boxplots show the median and interquartile range. P values were calculated using two-sided cell-level Wilcoxon rank-sum tests (n=12 mice pooled into two replicates). (E) Absolute counts of MHC-II^high^ BAMs (CD45^hi^ CD3^−^ CD19^−^ NK1.1^−^ CD11b^+^ Ly6C^−^ CD206^+^) in the brain as determined by flow cytometry 2.5m after intrathecal cell infusion (UT young, m.uPAR-m.28z young, UT old: n=10 mice; m.uPAR-m.28z old: n=8 mice). (F) Absolute counts of uPAR^+^ MHC-II^high^ BAMs (CD45^hi^ CD3^−^ CD19^−^ NK1.1^−^ CD11b^+^ Ly6C^−^ CD206^+^) in the brain as determined by flow cytometry 2.5m after intrathecal cell infusion (UT young, m.uPAR-m.28z young, UT old: n=10 mice; m.uPAR-m.28z old: n=8 mice). (G) Split-violin plot of the mean LogNormalized expression of the available genes from the myeloid inflammatory signature^42^ in BAMs from old UT and old m.uPAR-m.28z mice 2.5 months after intrathecal infusion. Boxplots show the median and interquartile range. P values were calculated using two-sided cell-level Wilcoxon rank-sum tests (old UT: n=6 mice pooled into two replicates; old m.uPAR-m.28z: n=4 mice pooled into two replicates). (H) Enrichr over-representation analysis^43^ of direction-separated genes differentially expressed between old UT and old m.uPAR-m.28z brains 2.5 months after intrathecal infusion. Odds ratio is shown on the x axis, nominal enrichment P value by color and Enrichr combined score by point size (old UT: n=6 mice pooled into two replicates; old m.uPAR-m.28z: n=4 mice pooled into two replicates). (I) Heatmap depicting the fold change in the protein levels of proinflammatory cytokines and chemokines in the brains of old UT or old m.uPAR-m.28z CAR T treated mice 20 days after intrathecal cell infusion (n = 5 mice per group). (J) Absolute counts of MHC-II^high^ BAMs (CD45^hi^ CD3^−^ CD19^−^ NK1.1^−^ CD11b^+^ Ly6C^−^ CD206^+^) in the brain as determined by flow cytometry 16m after intrathecal cell infusion (UT: n=12 mice; m.uPAR-m.28z: n=11 mice). (K) Absolute counts of uPAR^+^ MHC-II^high^ BAMs (CD45^hi^ CD3^−^ CD19^−^ NK1.1^−^ CD11b^+^ Ly6C^−^ CD206^+^) in the brain as determined by flow cytometry 16m after intrathecal cell infusion (UT: n=12 mice; m.uPAR-m.28z: n=11 mice). (L) Number of uPAR+ BAMs in the brain as determined by immunofluorescence staining of uPAR and CD206 6m after intrathecal cell infusion in the PS19 mice (UT: n=6 mice; m.uPAR-m.28z: n=7 mice). (M) Percentage of AT8 Tau positive area of total histology data as determined by immunohistochemistry 6m after intrathecal cell infusion in the PS19 mice (UT: n=7 mice; m.uPAR-m.28z: n=7 mice). (N-P) Analysis of the adult human brain atlas snRNA-seq dataset^19^ (888,263 nuclei from four donors); published cell annotations and the published UMAP embedding were retained. (N) UMAP-based binned density-difference map of the 888,263 retained nuclei. PLAUR^+^ nuclei were defined by an observed raw PLAUR count greater than zero. PLAUR^+^ and PLAUR^−^ nuclei were counted in a common two-dimensional grid on the published UMAP embedding; each grid was normalized by the total number of nuclei in that group and Gaussian-smoothed, after which PLAUR^−^ density was subtracted from PLAUR^+^ density. Red indicates relative enrichment of PLAUR^+^ nuclei and blue indicates relative enrichment of PLAUR^−^ nuclei; labels denote the 11 published non-neuronal cell classes. (O) Enrichr over-representation analysis^43^ of direction-separated genes differentially expressed between observed PLAUR^+^ and PLAUR^−^ nuclei. Odds ratio is shown on the x axis, nominal enrichment P value by color and Enrichr combined score by point size. (P) Split-violin plot of the myeloid inflammatory signature^42^ scored with Seurat’s *AddModuleScore* in observed PLAUR^−^ and PLAUR^+^ central nervous system macrophages. Boxplots show the median and interquartile range. P values were calculated using two-sided cell-level Wilcoxon rank-sum tests (PLAUR-: n=85,853 cells; PLAUR+: n=5,985 cells). (Q-R) Analysis of the FreshMG scRNA-seq dataset^20^. The age comparison includes one young donor (3,972 cells) and 11 old donors (38,903 cells). (Q) Split-violin plot of ALRA^41^-imputed PLAUR expression in PVMs from young (20-29 years) and old (90+ years) FreshMG brains. Boxplots show the median and interquartile range. P values were calculated using two-sided cell-level Wilcoxon rank-sum tests. (R) Split-violin plot of the myeloid inflammatory signature^42^ scored with Seurat’s *AddModuleScore* in observed PLAUR^−^ and PLAUR^+^ PVMs from old FreshMG donors. Boxplots show the median and interquartile range. P values were calculated using two-sided cell-level Wilcoxon rank-sum tests (PLAUR-: n=2,464 cells; PLAUR+: n=858 cells). (S-T) Age-matched PsychAD snRNA-seq analysis^20^ comprising 35,175 control nuclei from 214 donors and 85,697 Alzheimer disease nuclei from 434 donors. (S) Split-violin plot of ALRA^41^-imputed PLAUR expression in control and Alzheimer disease PVMs from the age-matched PsychAD cohort. Boxplots show the median and interquartile range. P values were calculated using two-sided cell-level Wilcoxon rank-sum tests (control: n=3,532 cells; Alzheimer disease: n=14,295 cells). (T) Split-violin plot of the myeloid inflammatory signature^42^ scored with Seurat’s *AddModuleScore* in observed PLAUR^−^ and PLAUR^+^ PVMs from Alzheimer disease donors. Boxplots show the median and interquartile range. P values were calculated using two-sided cell-level Wilcoxon rank-sum tests (PLAUR-: n=11,690 cells; PLAUR+: n=2,605 cells). Data are the mean ± s.e.m. (E-F) and (J-M). Statistical analysis was performed using two-tailed unpaired Student’s t-test (E-F) and (J-M); two-sided cell-level Wilcoxon rank-sum tests for the normalized-expression-average (D, G), AddModuleScore (P, R, T) and ALRA-expression (Q, S) comparisons; and Enrichr over-representation analysis for panels C, H and O. For the scRNA-seq enrichment panels, genes detected in at least 10% of either group with nominal P<0.05 and absolute average log2 fold change >0.5 were separated by direction for analysis. Data (B-D) and (G-H) represent one independent experiment or (E-F) and (J-M) two independent experiments.

To analyze how this population was modulated by intrathecal uPAR CAR T cells, we performed flow cytometry and histological analysis of mouse brains 2.5 months after treatment (a timepoint correlating with the observed cognitive improvements **(Figure 1D-F)**, and detected a significant decrease in the age-related accumulation of uPAR^+^ BAMs compared with controls **(Figure 2E-F and S2F)**. scRNAseq analysis of these brains at this time point, also revealed that reduction of uPAR^+^ BAMs significantly reduced the inflammatory nature of the remaining BAMs, resulting in an overall decrease of brain neuroinflammation as observed by downregulation of inflammatory terms through pathway analysis of differentially expressed genes in aged treated compared to control mice **(Figure 2G-H and S2G-J)**. Moreover, we also found proinflammatory cytokine and chemokine protein levels were decreased in the brains of uPAR CAR T treated aged mice **(Figure 2I)**.

Similar reduction of uPAR^+^ BAMs and decreased neuroinflammation was also observed in the brains of aged animals treated with a single CAR T infusion in their youth, 16 months after infusion **(Figure 2J-K and S2K-L)**. Analogously, uPAR CAR T cell treatment of PS19 mice also resulted in a significant reduction in the brain levels of uPAR^+^ BAMs, proinflammatory cytokines and chemokines and tau accumulation **(Figure 2L-M and S2M-O)**.

To understand whether these findings could be translatable to humans, we first analyzed the expression of *PLAUR*, the gene encoding uPAR, in snRNAseq data from the human brain atlas^19^. Across the identified human brain cell types, central nervous system macrophages exhibited the highest expression of *PLAUR,* and *PLAUR^+^* cells were significantly enriched in the expression of genes associated with inflammatory pathways **(Figure 2N-P)**. To examine in further detail how *PLAUR* expression changes in myeloid populations in human brains during aging and disease, we analyzed human prefrontal cortex scRNAseq and snRNAseq datasets enriched for myeloid populations^20^. Within human myeloid cells, and specifically, within perivascular macrophages (PVMs) (the closest human equivalent to BAMs^21^), *PLAUR* expression significantly increased during aging and Alzheimer’s disease, and, as in murine brains, identified a subset of highly inflammatory PVMs **(Figure 2Q-T and S2P-S)**.

Collectively, these data identify uPAR^+^ BAMs as a highly inflammatory subset that increases in mouse and human aged brains and in neurodegeneration. Moreover, this subset can be targeted through intrathecal administration of uPAR CAR T cells resulting in decreased neuroinflammation.

### Intravenous administration of CAR T cells phenocopies intrathecal treatment

Recent studies have suggested disruption of CNS homeostasis, such as during infections, acute brain injury, stroke, under experimental irradiation, or pharmacological-induced depletion, can result in BAMs being partially replaced by monocyte-derived cells from the bone marrow^9–12^. Whether this process also occurs during aging and how it contributes to age-associated neuroinflammation remains unclear.

To investigate whether peripheral sources contribute to age-associated inflammatory uPAR^+^ BAMs, we examined the cognitive function of aged mice after intravenous administration of uPAR CAR T cells, which does not result in significant brain infiltration **(Fig. 1B)**. Surprisingly, intravenous uPAR CAR T cell administration also led to improvements in cognitive performance in old mice, as well as to a reduction in the number of uPAR^+^ BAMs and a downregulation in the protein levels of proinflammatory cytokines and chemokines in the brains of these animals **(Figure 3A-F and S3A-G)**. However, while intrathecal administration elicited cognitive improvements after 2.5 months, intravenous treatment took 6 months **(Figure 3A-C and S3A-C),** consistent, possibly, with potential targeting of bone marrow progenitors whose progeny traffic to the brain.

**Figure 3.**
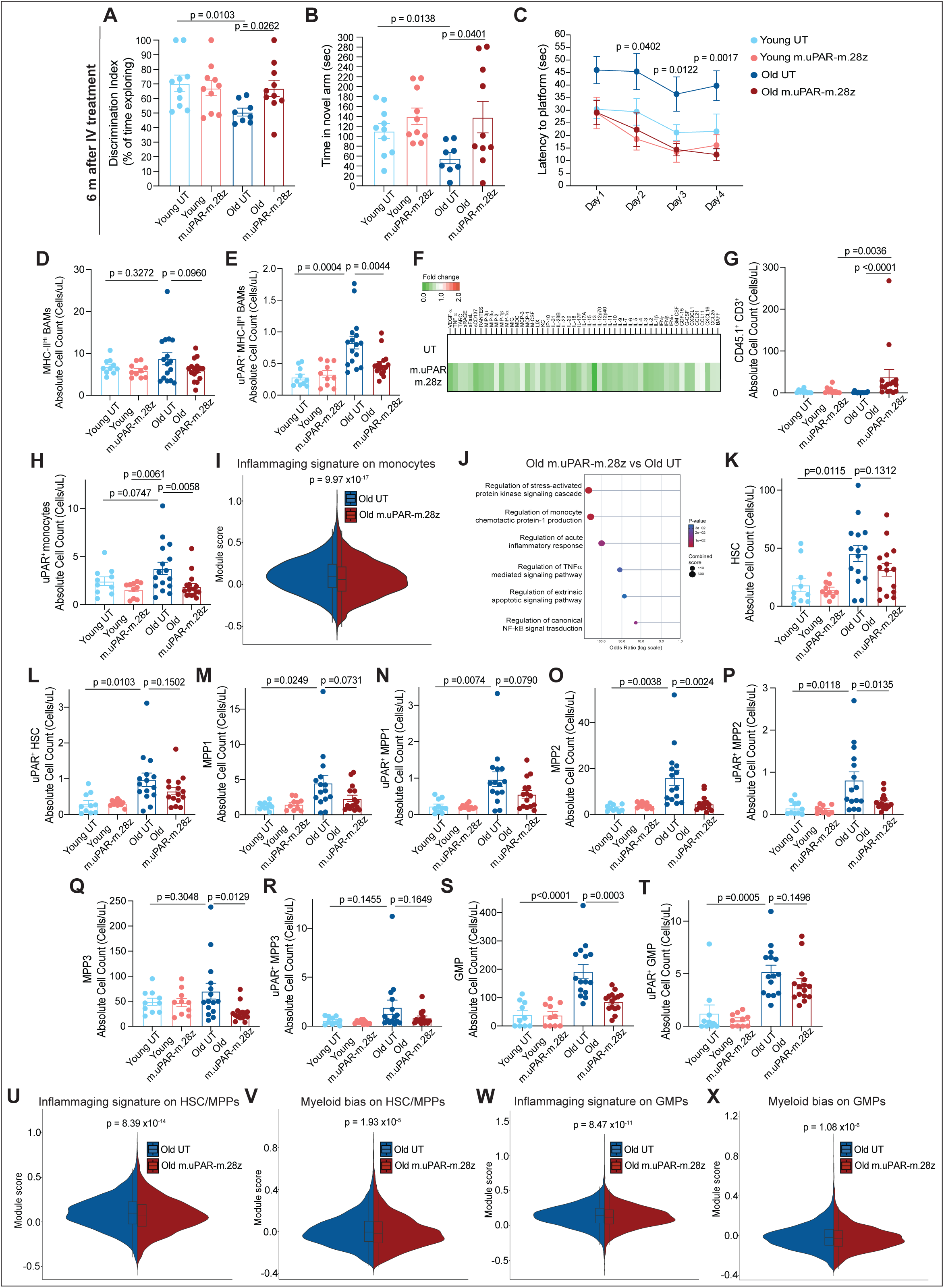
Intravenous uPAR CAR T cells phenocopy the brain effects of intrathecal uPAR CAR T cells. (A) Discrimination Index in the novel object recognition task 6m after intravenous cell infusion. (UT young, m.uPAR-m.28z young: n=10 mice per group, UT old: n=8 mice; m.uPAR-m.28z old: n=10 mice). (B) Time spent in novel arm in the Y-maze spatial memory task 6m after intravenous cell infusion. (UT young, m.uPAR-m.28z young: n=10 mice per group, UT old: n=8 mice; m.uPAR-m.28z old: n=10 mice). (C) Latency to the platform in the Morris water maze test 6m after intravenous cell infusion. (n=6 mice per group). (D) Absolute counts of MHC-II^high^ BAMs (CD45^hi^ CD3^−^ CD19^−^ NK1.1^−^ CD11b^+^ Ly6C^−^ CD206^+^) in the brain as determined by flow cytometry 6m after intravenous cell infusion (UT young, m.uPAR-m.28z young, n=10 mice; UT old, m.uPAR-m.28z old: n=16 mice). (E) Absolute counts of uPAR^+^ MHC-II^high^ BAMs (CD45^hi^ CD3^−^ CD19^−^ NK1.1^−^ CD11b^+^ Ly6C^−^ CD206^+^) in the brain as determined by flow cytometry 6m after intravenous cell infusion (UT young, m.uPAR-m.28z young, n=10 mice; UT old, m.uPAR-m.28z old: n=16 mice). (F) Heatmap depicting the fold change in the protein levels of proinflammatory cytokines and chemokines in the brains of old UT or old m.uPAR-m.28z CAR T treated mice 6m after intravenous cell infusion (n = 4 mice per group). (G) Absolute counts of CD45.1 and CD3 double positive cells in the bone marrow as determined by flow cytometry 6 months after intravenous cell infusion. (UT young, m.uPAR-m.28z young: n=10 mice; UT old: n=16 mice, m.uPAR-m.28z old: n=15 mice). (H) Absolute counts of uPAR positive monocytes (CD45+ CD3- CD19- NK1.1- CD11b+ Ly6C+ Ly6G-) in the bone marrow as determined by flow cytometry 6 months after intravenous cell infusion. (UT young, m.uPAR-m.28z young: n=10 mice; UT old and m.uPAR-m.28z old: n=16 mice). (I) Split-violin plot of the Chambers et al. inflammaging signature^44^ scored with Seurat’s *AddModuleScore* in bone-marrow monocytes from old UT and old m.uPAR-m.28z mice 6 months after intravenous infusion. Boxplots show the median and interquartile range. P values were calculated using two-sided cell-level Wilcoxon rank-sum tests (n=4 mice per group pooled into two replicates). (J) Enrichr over-representation analysis^43^ of direction-separated genes differentially expressed between old UT and old m.uPAR-m.28z bone marrow 6 months after intravenous infusion. Odds ratio is shown on the x axis, nominal enrichment P value by color and Enrichr combined score by point size (n=4 mice per group pooled into two replicates). (K) Absolute counts of HSCs (Sca-1+ cKit+ CD150+ CD48- CD135- CD34-) in the bone marrow as determined by flow cytometry 6 months after intravenous cell infusion. (UT young, m.uPAR-m.28z young: n=10 mice; UT old and m.uPAR-m.28z old: n=15 mice). (L) Absolute counts of uPAR^+^ HSCs (Sca-1+ cKit+ CD150+ CD48- CD135- CD34-) in the bone marrow as determined by flow cytometry 6 months after intravenous cell infusion. (UT young, m.uPAR-m.28z young: n=10 mice; UT old and m.uPAR-m.28z old: n=15 mice). (M) Absolute counts of MPP1s (Sca-1+ cKit+ CD150+ CD48- CD135- CD34+) in the bone marrow as determined by flow cytometry 6 months after intravenous cell infusion. (UT young, m.uPAR-m.28z young: n=10 mice; UT old and m.uPAR-m.28z old: n=15 mice). (N) Absolute counts of uPAR^+^ MPP1s (Sca-1+ cKit+ CD150+ CD48- CD135- CD34+) in the bone marrow as determined by flow cytometry 6 months after intravenous cell infusion. (UT young, m.uPAR-m.28z young: n=10 mice; UT old and m.uPAR-m.28z old: n=15 mice). (O) Absolute counts of MPP2s (Sca-1+ cKit+ CD150+ CD48+) in the bone marrow as determined by flow cytometry 6 months after intravenous cell infusion. (UT young, m.uPAR-m.28z young: n=10 mice; UT old and m.uPAR-m.28z old: n=15 mice). (P) Absolute counts of uPAR^+^ MPP2s (Sca-1+ cKit+ CD150+ CD48+) in the bone marrow as determined by flow cytometry 6 months after intravenous cell infusion. (UT young, m.uPAR-m.28z young: n=10 mice; UT old and m.uPAR-m.28z old: n=15 mice). (Q) Absolute counts of MPP3s (Sca-1+ cKit+ CD150- CD48+ CD135-) in the bone marrow as determined by flow cytometry 6 months after intravenous cell infusion. (UT young, m.uPAR-m.28z young: n=10 mice; UT old and m.uPAR-m.28z old: n=15 mice). (R) Absolute counts of uPAR^+^ MPP3s (Sca-1+ cKit+ CD150- CD48+ CD135-) in the bone marrow as determined by flow cytometry 6 months after intravenous cell infusion. (UT young, m.uPAR-m.28z young: n=10 mice; UT old and m.uPAR-m.28z old: n=15 mice). (S) Absolute counts of GMPs (Sca-1-cKit+ CD16/32+ CD34+) in the bone marrow as determined by flow cytometry 6 months after intravenous cell infusion. (UT young, m.uPAR-m.28z young: n=10 mice; UT old and m.uPAR-m.28z old: n=15 mice). (T) Absolute counts of uPAR^+^ GMPs (Sca-1-cKit+ CD16/32+ CD34+) in the bone marrow as determined by flow cytometry 6 months after intravenous cell infusion. (UT young, m.uPAR-m.28z young: n=10 mice; UT old and m.uPAR-m.28z old: n=15 mice). (U) Split-violin plot of the Chambers et al. inflammaging signature^44^ scored with Seurat’s *AddModuleScore* in HSC/MPP populations from old UT and old m.uPAR-m.28z marrow. Boxplots show the median and interquartile range. P values were calculated using two-sided cell-level Wilcoxon rank-sum tests (n=4 mice per group pooled into two replicates). (V) Split-violin plot of the Kovtonyuk et al. myeloid-bias signature^45^ scored with Seurat’s *AddModuleScore* in HSC/MPP populations from old UT and old m.uPAR-m.28z marrow. Boxplots show the median and interquartile range. P values were calculated using two-sided cell-level Wilcoxon rank-sum tests (n=4 mice per group pooled into two replicates). (W) Split-violin plot of the Chambers et al. inflammaging signature^44^ scored with Seurat’s *AddModuleScore* in GMPs from old UT and old m.uPAR-m.28z marrow. Boxplots show the median and interquartile range. P values were calculated using two-sided cell-level Wilcoxon rank-sum tests (n=4 mice per group pooled into two replicates). (X) Split-violin plot of the Kovtonyuk et al. myeloid-bias signature^45^ scored with Seurat’s *AddModuleScore* in GMPs from old UT and old m.uPAR-m.28z marrow. Boxplots show the median and interquartile range. P values were calculated using two-sided cell-level Wilcoxon rank-sum tests (n=4 mice per group pooled into two replicates). Data are the mean ± s.e.m. (A-E), (G-H) and (K-T). Statistical analysis was performed using two-tailed unpaired Student’s t-test (A-E) and (K-T); Mann-Whitney test (G); Kruskal-Wallis test (H); two-sided cell-level Wilcoxon rank-sum tests for the AddModuleScore (I,U-X); and Enrichr over-representation analysis for (J). For (J), genes detected in at least 10% of either group with nominal P<0.05 and absolute average log2 fold change >0.5 were separated by direction for analysis. Data (A-E), (G-H) and (K-T) represent two independent experiments or (F), (I-J),(U-X) one independent experiment.

Analysis of the peripheral bone marrow revealed a significant presence of uPAR CAR T cells in aged mice, which displayed an effector phenotype **(Figure 3G and S3H-I)**. uPAR CAR T cell treatment resulted in a significant decrease in the number of uPAR^+^ monocytes in the bone marrow of aged mice, comparable to the levels present in young animals **(Figure 3H and S3J)**. Furthermore, scRNAseq analysis showed significant downregulation of overall inflammation in the bone marrow, with the remaining monocytes being significantly less inflammatory than those of aged UT treated animals **(Figure 3I-J and S3K-N)**.

During aging, the mouse bone marrow undergoes an expansion of hematopoietic stem cells (HSCs) and multipotent progenitors (MPP1, MPP2, MPP3) that are inflammatory and biased towards the myeloid lineage^22,23^. Intravenous treatment with the uPAR CAR T cells reduced the numbers of uPAR^+^ HSCs, MPP1, MPP2, MPP3 and GMP populations, which increased with aging. Additionally, uPAR CAR T cell treatment resulted in a reduction in the age-related increase in total numbers of these populations, suggesting that the remaining uPAR^−^ progenitors were “rejuvenated” **(Figure 3K-T)**. Indeed, scRNAseq of bone marrow cells at this time point showed that treatment with the uPAR CAR T cells reduced the expression of age-related inflammatory and myeloid bias programs in these progenitor populations **(Figure 3U-X)**.

Taken together, these data identify an age-associated increase in uPAR^+^ myeloid progenitors in the bone marrow, whose targeting by intravenous delivery of CAR T cells reduces bone marrow inflammation and myeloid bias. Notably, these progenitor changes in the bone marrow correlate with concomitant decreases in inflammatory uPAR+ BAMs in the brain and improvements in cognition.

### CAR T cell targeting of bone marrow inflammatory myeloid progenitors is sufficient to improve age-associated cognitive impairment

To better understand whether the effects of intravenous uPAR CAR T cells are the result of directly targeting uPAR^+^ myeloid progenitors or if the elimination of uPAR^+^ cells in other tissues is also necessary, we performed a bone marrow transplant experiment. UT or uPAR CAR T cells were generated from CD45.1 mice and infused intravenously into aged (18 months old) CD45.2 animals. 6 weeks after administration, we isolated 200,000 myeloid-biased progenitors from the bone marrow of these animals as well as from a young control and transplanted them into aged CD45.1 mice **(Figure 4A)**. Engraftment numbers of total exogenous CD45.2 cells in the bone marrow were comparable among the three groups, as were the total numbers of exogenous CD45.2 myeloid cells that migrated into the aged brains, confirming that myeloid progenitors in the bone marrow are a source of BAMs in aging **(Figure 4B-C and S4A-D)**.

**Figure 4.**
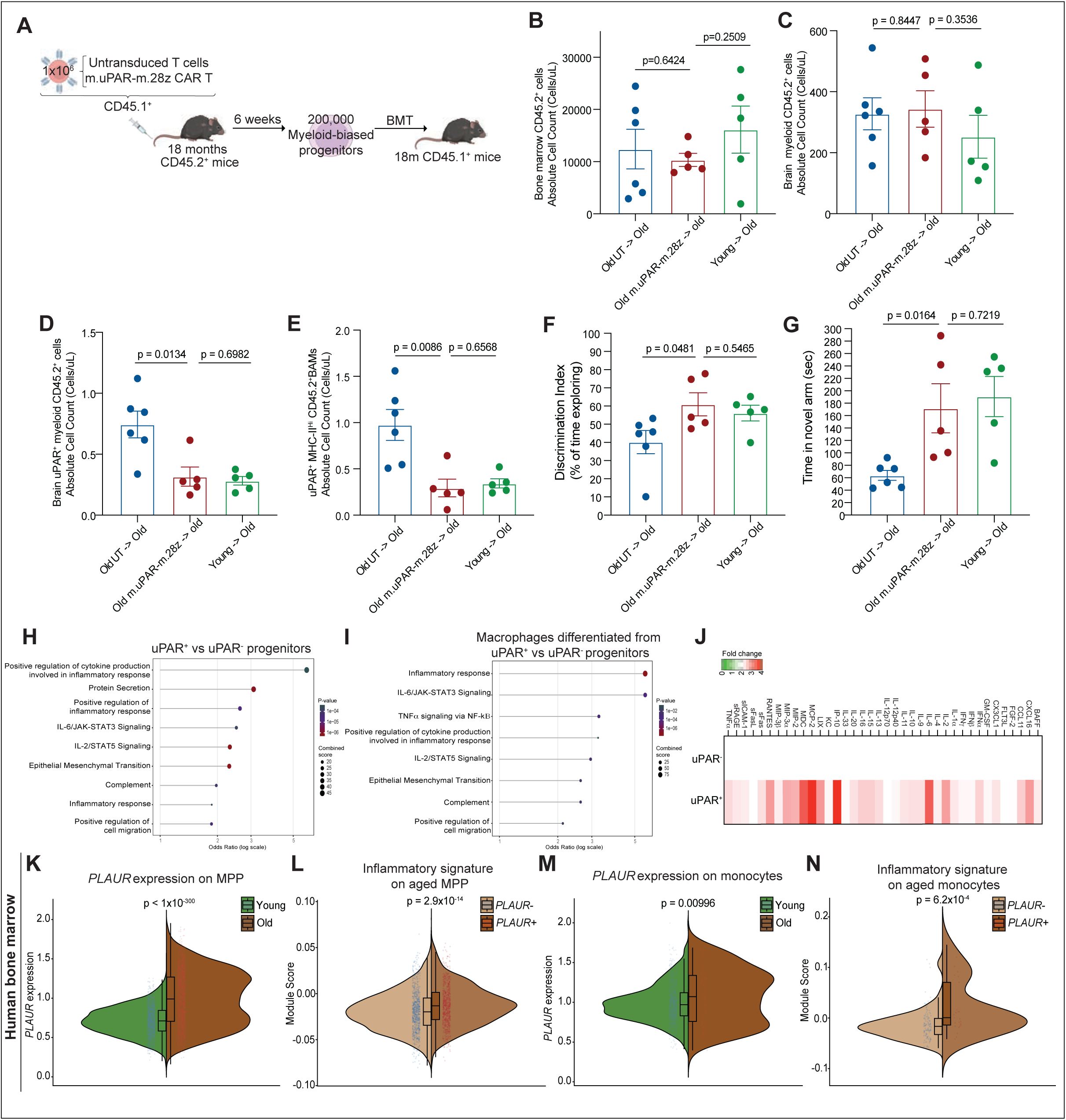
uPAR CAR T cells targeting of bone marrow progenitors is sufficient to improve age-associated cognitive impairment. (A) Schematic of experimental design for (B-G): Old (18 months) CD45.2^+^mice were intravenously infused with either 1×10^6^ untransduced T cells (UT) or m.uPAR-m.28z CAR T cells generated from CD45.1^+^ mice. Six weeks later, hematopoietic stem and progenitor cells were isolated from the bone marrow, biased towards myeloid lineage overnight by culture in M-CSF and IL6 and transplanted into old (17-22 months CD45.1^+^ mice that had been preconditioned with busulfan (20 mg kg^−1^ for 3 consecutive days). Transplanted mice underwent behavioral testing 3 months later and flow cytometry of bone marrow and brain 5 months later. (B) Absolute counts of CD45.2 positive cells in the bone marrow as determined by flow cytometry 5 months after bone marrow transplant (Old UT to old: n=6 mice, Young to old and old m.uPAR-m.28z to old: n=5). (C) Absolute counts of total CD45.2 CD11b positive cells in the brain as determined by flow cytometry 5 months after bone marrow transplant (Old UT to old: n=6 mice, Young to old and old m.uPAR-m.28z to old: n=5). (D) Absolute counts of total uPAR^+^ CD45.2 CD11b positive cells in the brain as determined by flow cytometry 5 months after bone marrow transplant (Old UT to old: n=6 mice, Young to old and old m.uPAR-m.28z to old: n=5). (E) Absolute counts of uPAR^+^ MHC-II^high^ CD45.2^+^ BAMs in the brain as determined by flow cytometry 5 months after bone marrow transplant (Old UT to old: n=6 mice, Young to old and old m.uPAR-m.28z to old: n=5). (F) Discrimination Index in the novel object recognition task 3 months after bone marrow transplant (Old UT to old: n=6 mice, Young to old and old m.uPAR-m.28z to old: n=5). (G) Time spent in novel arm in the Y-maze spatial memory task 3 months after bone marrow transplant (Old UT to old: n=6 mice, Young to old and old m.uPAR-m.28z to old: n=5). (H) Pathway analysis using enrichR comparing differentially expressed genes between sorted uPAR+ and uPAR- lineage- bone marrow cells. Size scale represents number of genes in each ontology, and color scale represents degree of significance (n=3 mice per group). (I) Pathway analysis using enrichR comparing differentially expressed genes between macrophages differentiated in vitro from uPAR^+^ and uPAR^−^ lineage^−^ bone marrow cells by addition of 25ng/ml of M-CSF for one week. Size scale represents number of genes in each ontology, and color scale represents degree of significance (n=3 mice per group). (J) Heatmap depicting the fold change in the protein levels of proinflammatory cytokines and chemokines between macrophages differentiated in vitro from uPAR^+^ and uPAR^−^ lineage^−^ bone marrow cells by addition of 25ng/ml of M-CSF for one week (n = 3 mice per group). (K) Split-violin plot of ALRA^41^-imputed PLAUR expression in healthy human bone-marrow MPP-like cells from Ainciburu et al^56^. Boxplots show the median and interquartile range. P values were calculated using two-sided cell-level Wilcoxon rank-sum tests (young: n=9,170 cells from five donors; elderly: n=5,780 cells from three donors). (L) Split-violin plot of the MSigDB Hallmark Inflammatory Response gene set^57^ scored with Seurat’s *AddModuleScore* in observed PLAUR^−^ and PLAUR^+^ MPP-like cells from elderly donors in Ainciburu et al.^56^. Boxplots show the median and interquartile range. P values were calculated using two-sided cell-level Wilcoxon rank-sum tests (PLAUR^−^: n=4,945 cells; PLAUR^+^: n=835 cells). (M) Split-violin plot of ALRA^41^-imputed PLAUR expression in healthy human bone-marrow monocytes from Ainciburu et al. ^56^. Boxplots show the median and interquartile range. P values were calculated using two-sided cell-level Wilcoxon rank-sum tests (young: n=717 cells from five donors; elderly: n=140 cells from three donors). (N) Split-violin plot of the MSigDB Hallmark Inflammatory Response gene set^57^ scored with Seurat’s *AddModuleScore* in observed PLAUR^−^ and PLAUR^+^ monocytes from elderly donors in Ainciburu et al. ^56^. Boxplots show the median and interquartile range. P values were calculated using two-sided cell-level Wilcoxon rank-sum tests (PLAUR^−^: n=115 cells; PLAUR^+^: n=25 cells). Data are the mean ± s.e.m. (B-G). Statistical analysis was performed using two-tailed unpaired Student’s t-test (B-G), and significance values depicted in the color scale represent adjusted p-values calculated using Fisher’s exact test (H-I) or two-tailed Wilcoxon rank-sum test (K-N). Data (B-N) represent one independent experiment.

However, the numbers of exogenous uPAR^+^ CD45.2^+^ monocytes in the bone marrow and uPAR^+^ CD45.2^+^ BAMs were significantly decreased in the aged mice that received the myeloid progenitors from aged uPAR CAR T treated animals compared with aged UT treated mice; this reduction was comparable to aged mice receiving myeloid progenitors from young mice **(Figure 4D-E and S4D-F)**. The reduction in uPAR^+^ BAMs coincided with cognitive improvements in recognition and spatial memory to levels similar to those obtained by transplanting myeloid progenitors from young mice **(Figure 4F-G and S4O-P)**. Collectively, these data suggest that targeting of uPAR^+^ bone marrow myeloid progenitors is sufficient to: 1) reduce the numbers of uPAR^+^ BAMs in the brain and 2) improve age-associated memory dysfunction.

Furthermore, within the endogenous immune system, the presence of uPAR CAR T cell-treated CD45.2 progenitors reduced myeloid bias in the endogenous bone marrow and uPAR expression in endogenous CD45.1 monocytes, resulting also in decreased uPAR^+^ CD45.1^+^ BAMs in the brain **(Figure S4G-N)**. This suggests that depletion of uPAR^+^ myeloid progenitors can also have cell extrinsic effects that rejuvenate the endogenous bone marrow compartment.

To further understand the nature of uPAR^+^ bone marrow progenitors, we performed bulk RNAseq on sorted uPAR^+^ and uPAR^−^ lineage^−^ cells cells from the bone marrow of 18 month old mice. Pathway analysis of differentially expressed genes revealed that uPAR+ Lin^−^ cells were highly enriched in inflammatory pathways compared to uPAR^−^ Lin^−^ counterparts **(Figure 4H)**. Furthermore, we found that in vitro differentiated macrophage progeny from uPAR^+^ progenitors had inherited a comparable inflammatory transcriptional state and secreted higher protein levels of proinflammatory cytokines and chemokines compared to macrophage progeny from uPAR^−^ progenitors **(Figure 4I-J)**.

Importantly, we observed that in scRNAseq from human bone marrow, *PLAUR* expression increased with age across HSCs, MPPs, GMP and monocytes, where it also identified inflammatory subsets **(Figure 4K-N and S4Q-T)**.

Collectively, these data suggest that uPAR surface expression on bone marrow myeloid progenitors marks a subset of inflammatory cells, which are conserved in human aging, and that transmit their inflamed state to their BAM progeny, which in turn drive age-associated cognitive impairment. Consequently, depleting uPAR^+^ cells with intravenous CAR T cells reduces inflammatory myeloid progenitors in the bone marrow and uPAR^+^ BAMs, decreasing neuroinflammation and improving cognitive performance.

## Discussion

Here we identify a subset of inflammatory BAMs as drivers of neuroinflammation and associated cognitive impairment, whose targeting through intrathecal CAR T cell delivery improves memory function in aged mice, and a mouse model of tauopathy, which also showed extended survival. Moreover, prophylactic administration of CAR T cells in young mice prevents cognitive impairment 16 months later. These inflammatory BAMs originate from a subset of bone marrow inflammatory myeloid progenitors that heritably transmit their inflammatory state to their progeny. Indeed, peripheral direct targeting of this myeloid progenitor population is sufficient to reduce neuroinflammation and improve memory function. The identification of uPAR as a surface marker of these progenitors, conserved in human aging, enables their precise therapeutic elimination, opening a new class of cell-targeting CAR T cell approaches for aging and neurodegeneration associated cognitive decline.

The identification of uPAR^+^ inflammatory BAMs as key mediators of neuroinflammation extends emerging evidence that BAMs acquire dysfunctional properties during brain aging and neurodegeneration such as Parkinson’s disease^24,25^. Of note, our therapeutic approach selectively targets the dysfunctional inflammatory uPAR^+^ subset, while preserving the broader BAM compartment, consistent with data that functionally intact BAMs can be neuroprotective in AD^26^.

The bone marrow ontogeny of this inflammatory BAM population during aging represents a previously unrecognized axis. Whether BAMs are replenished from bone marrow-derived monocytes remains debated: while a previous study has shown minimal bone marrow replacement of total BAMs in an amyloid-β AD model^27^, others have shown that monocyte-derived BAMs can replenish the BAM niche, particularly of the dural, leptomeninges and perivascular spaces especially under conditions of injury and disease^9–11,28^. Whether the progeny of bone marrow myeloid progenitors contribute to the BAM pool during aging and whether their expression profile contributes to neuroinflammation and cognitive decline had not been addressed. Our data provide evidence that inflammatory uPAR^+^ myeloid progenitors accumulate with age and give rise to inflammatory BAMs, establishing a causal link between bone marrow progenitor state and BAM-mediated neuroinflammation in aging. A recent study reported impaired myelopoiesis as a downstream consequence of amyloid β pathology^29^, our data show that inflammatory uPAR^+^ progenitor accumulation drives upstream neuroinflammation, suggesting these represent distinct but potentially interacting mechanisms.

The identification of targetable surface markers on inflammatory myeloid progenitors expands and distinguishes our work from prior descriptions of inflammatory hematopoietic states^30,31^. The broader normalization of the aged bone marrow, including a non-cell autonomous reduction in the myeloid bias of endogenous progenitors following transplantation of CAR T treated progenitors, may potentially reflect paracrine anti-inflammatory remodeling of the bone marrow niche, although contribution from reduced niche competition cannot be excluded. Of note, uPAR has been shown to play homeostatic roles in young HSCs by regulating mobilization^32^, suggesting its age-associated upregulation may reflect a hijacking of a physiological niche signal by the inflammatory aging microenvironment.

Our findings show that only a single peripheral intravenous CAR T cell administration is needed for durable improvements in cognitive performance, contrasting with current neurodegeneration therapies that require repeated dosing^2,3^. While further work is needed to examine the translatability of our findings, *PLAUR* is upregulated in human inflammatory perivascular macrophages in aging and AD brain datasets and in human bone marrow progenitors during aging. Interestingly, genetic studies have identified associations between *PLAUR* and *PLAU* pathway variants and increased AD risk^33,34^. Moreover, epidemiological evidence linking repeated infections to increased AD risk^35^ is consistent with our model, as inflammatory challenges alter the expression profile in hematopoietic progenitors that could amplify the bone marrow-BAM inflammatory axis we describe.

Collectively, our findings reframe the aging bone marrow as a peripheral driver of aging-associated neuroinflammation and cognitive impairment, and establish peripheral CAR T cell mediated resetting of this axis as a potential therapeutic and preventive strategy in neurodegeneration.

### Limitations of the study

While our data shows that elimination of uPAR^+^ BAMs reduces neuroinflammation, non-cell autonomous mechanisms whereby uPAR^+^ BAMs interact with other brain resident cell types (such as microglia, astrocytes and infiltrating immune cells) to collectively sustain neuroinflammation remain unexplored. Future work dissecting these interactions will be important to fully elucidate how BAM dysfunction propagates neuroinflammation and neurodegeneration in the aging brain.

Finally, given the existence of channels that connect the skull bone marrow to the meningeal spaces^28^, it is possible that intrathecal CAR T cell delivery partially exerts its effects through targeting of uPAR^+^ myeloid progenitors in the skull bone marrow in addition to the direct elimination of brain resident BAMs, which would further underscore the therapeutic relevance of peripheral targeting of the bone marrow-BAM axis in aging.

## Resource availability

### Lead contact

Requests for resources and reagents should be directed to the lead contact, Dr. Corina Amor.

## Acknowledgements

We would like to thank G.Alderton for editing the manuscript and L.Van Aelst and L.Cheadle for helpful discussions and feedback. We thank Cold Spring Harbor laboratory (CSHL) Cancer Center Shared Resources (Animal Facility, Flow Cytometry, Bioinformatics, Next Gen Sequencing, Microscopy, and Histology Core Facilities) supported by NCI Cancer Center Support grant 5P30CA045508. Special thanks to E. Earl, B. Rose and A. Bjertnes for outstanding animal care. Work in the laboratory of C.A. was supported by the National Institutes of Health Common Fund grant 1DP5OD033055 and National Institute on Aging grant 1R01AG082800, the Pershing Square MIND Prize and the Howard Hughes Medical Institute.

## Author contributions

A.S.H., J.A.R., G.G., I.F.M. and S.P. designed, performed and analyzed experiments. J.A.R. and R.U. performed scRNAseq analysis. B.O., S.L., W.Y.S., A.A., J.G., M.S., J.H., performed experiments. C.R.V. supervised experiments. C.A. conceived the project, acquired funding, designed, and supervised experiments, and wrote the paper with assistance from all authors. All authors read and approved the paper.

## Declaration of generative AI and AI-assisted technologies in the manuscript preparation process

During the preparation of this work the authors used Claude Sonnet 4.6 in order to edit the text. The authors reviewed and edited the content as needed and take full responsibility for the content of the published article.

## STAR Methods

### Key resources table

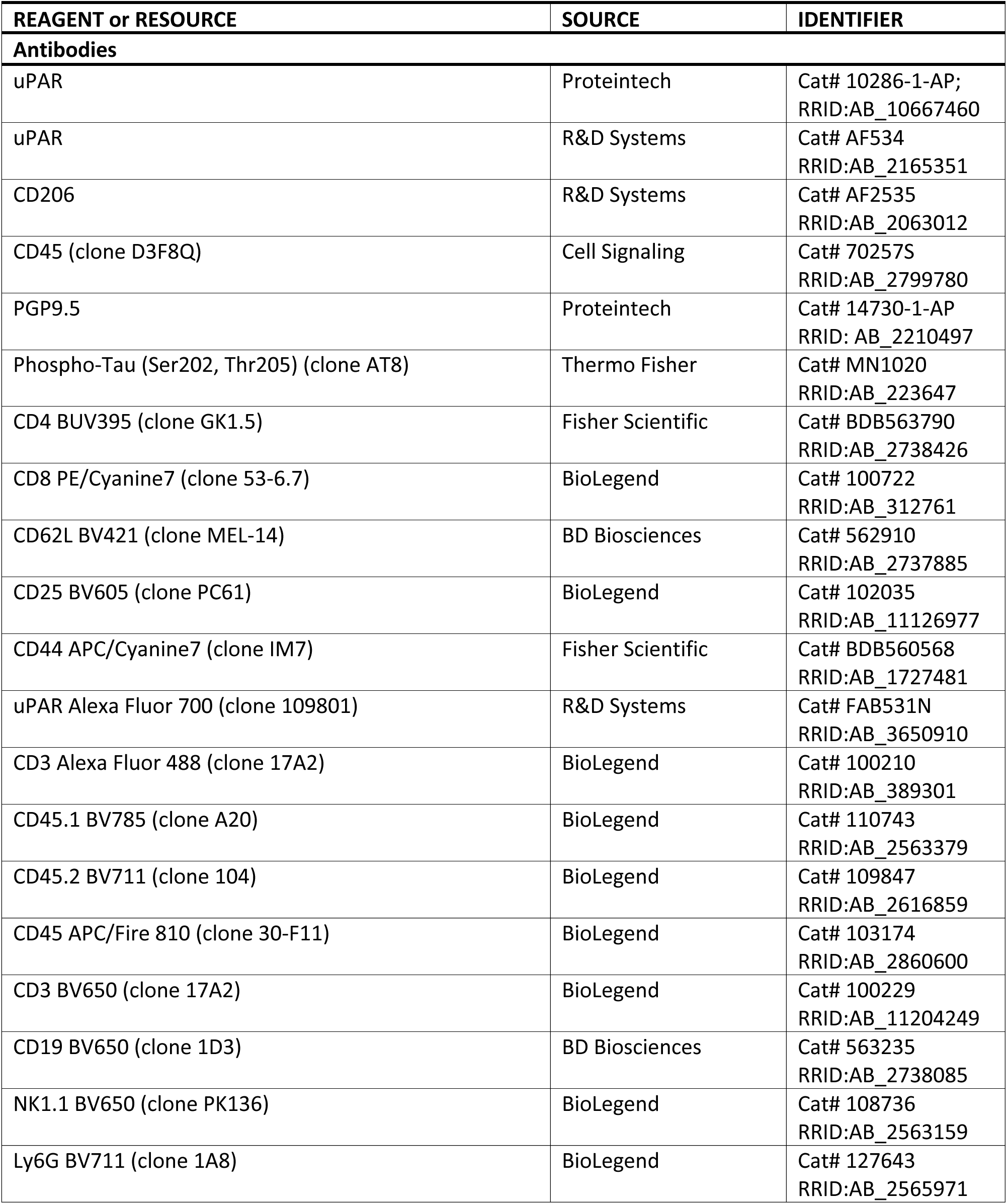
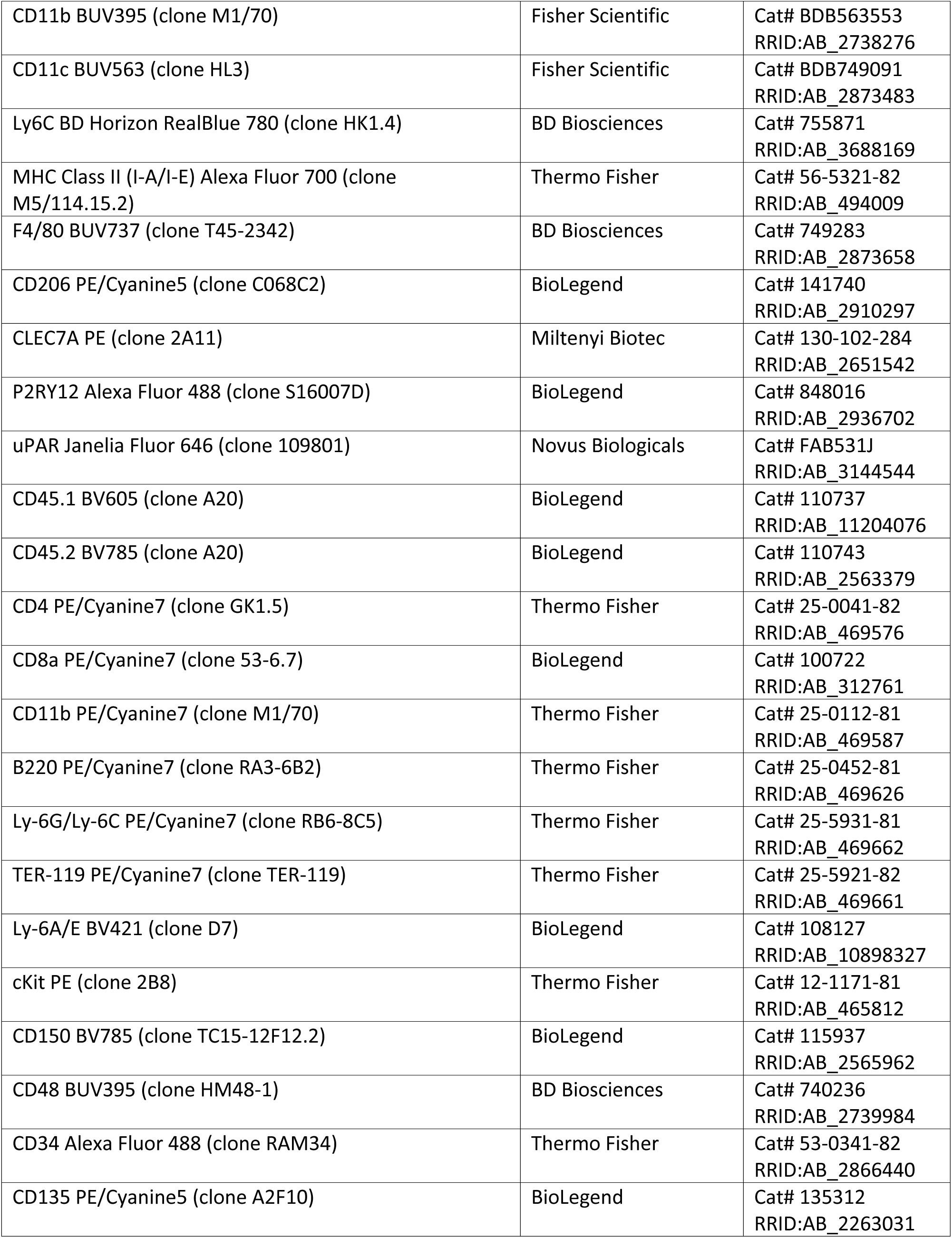
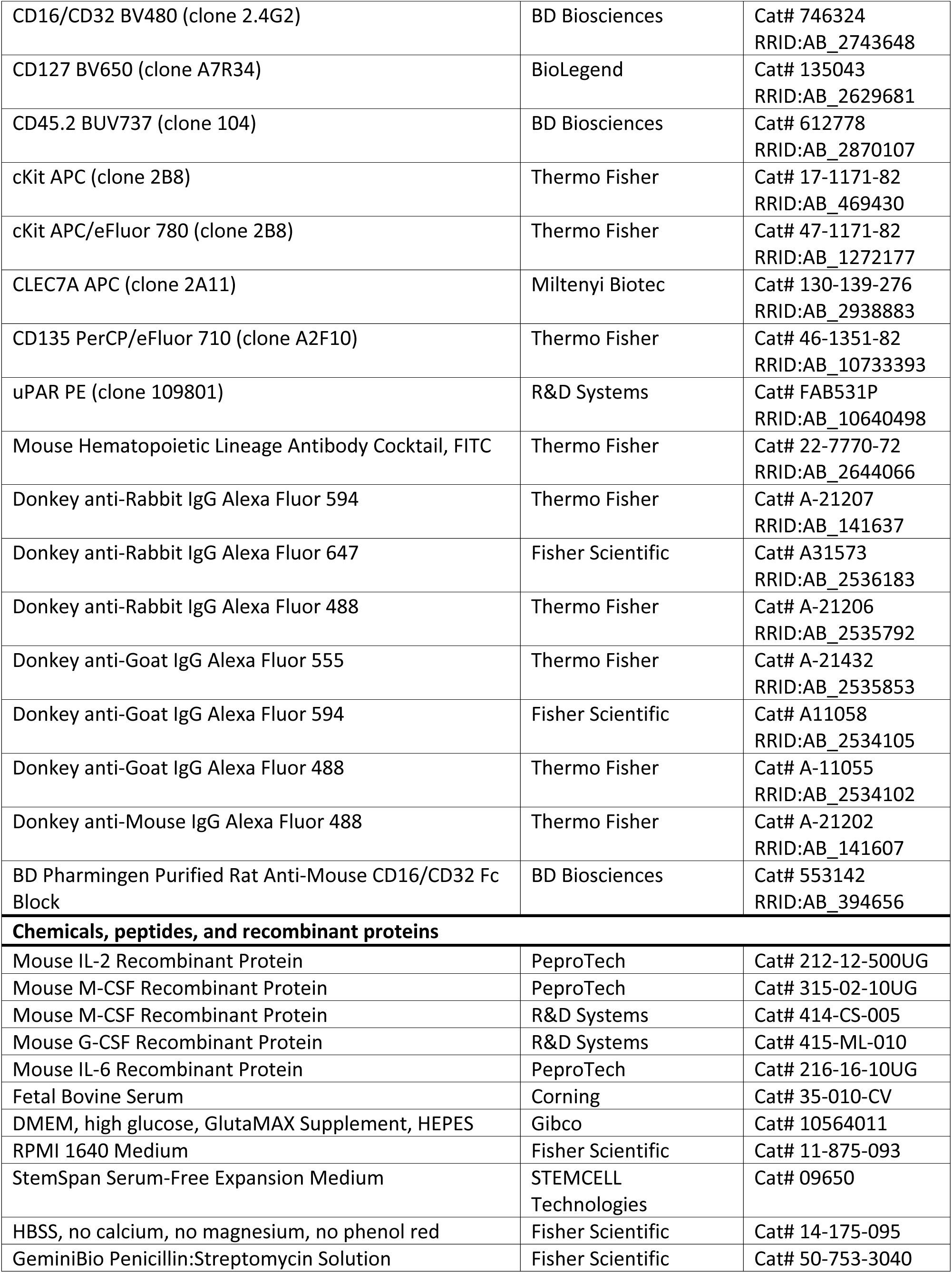
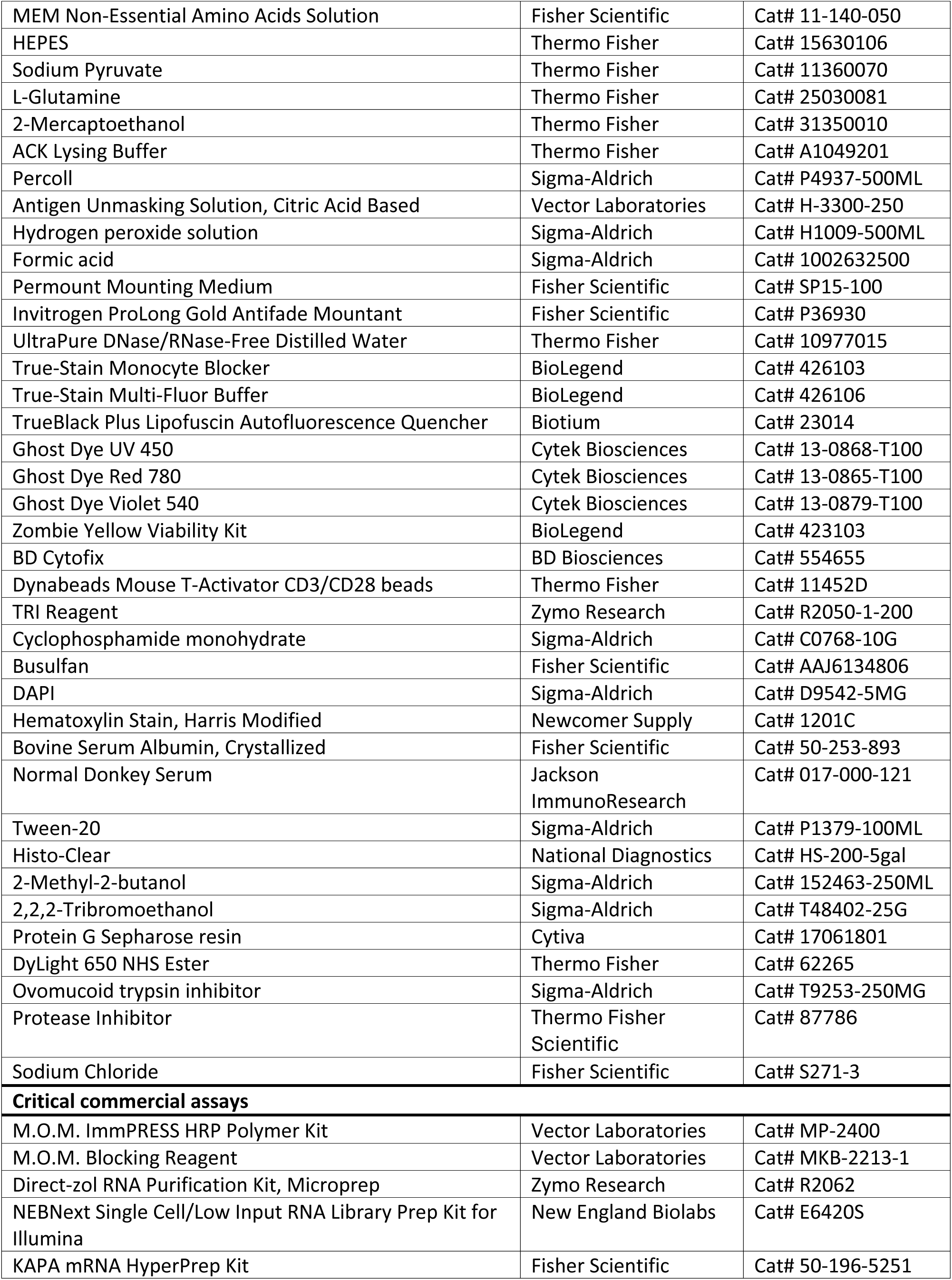
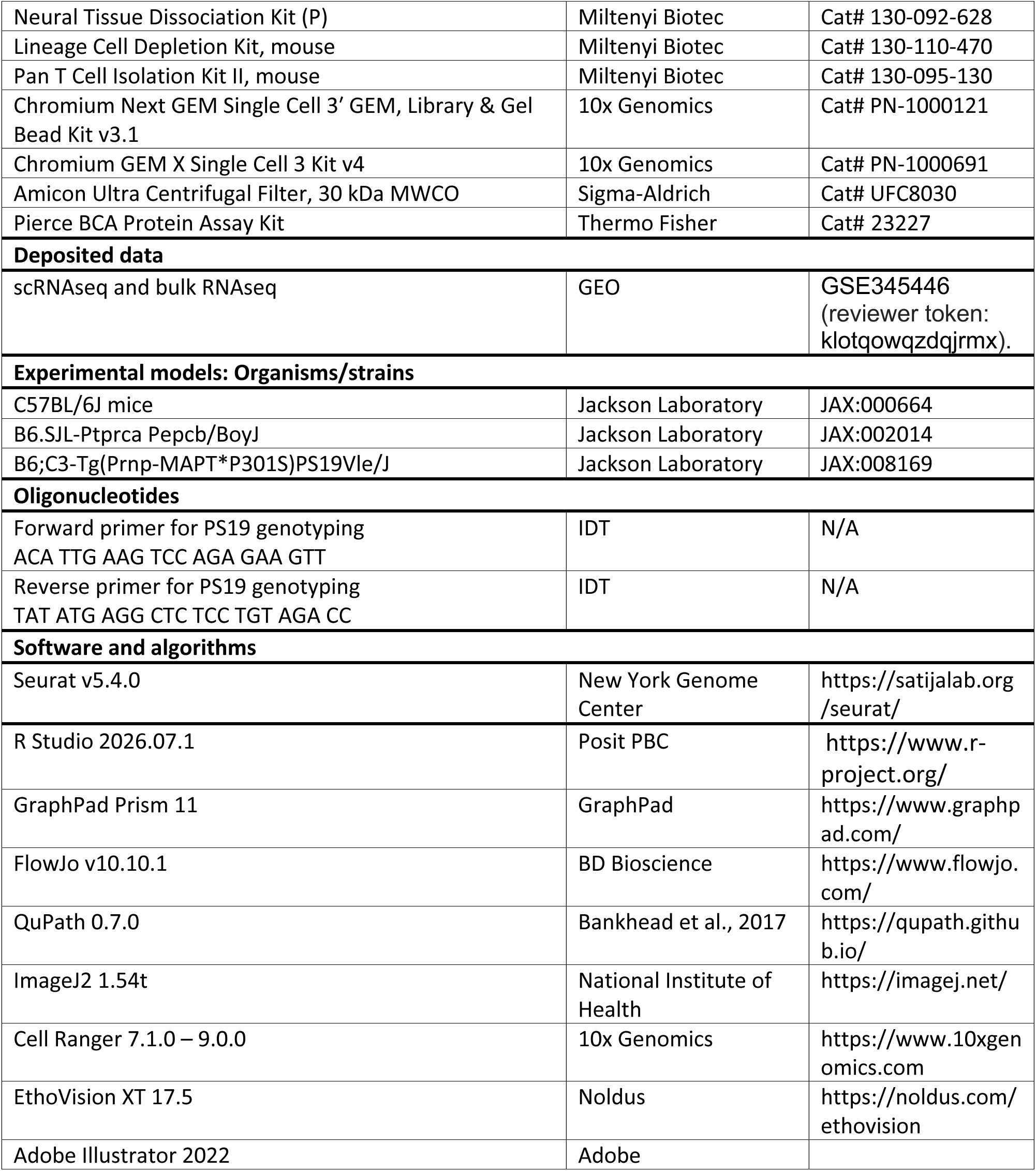

### Mouse strains and housing

C57BL/6J mice (JAX:000664), B6.SJL-Ptprca Pepcb/BoyJ mice (JAX:002014), and B6;C3-Tg(Prnp-MAPT*P301S)PS19Vle/J mice (JAX:008169) were purchased from The Jackson Laboratory. C57BL/6J mice were considered young at 8 to 12 weeks of age and aged at 72 to 76 weeks of age when treated with CAR T cells either intravenously (1 × 10⁶ cells per 200 µL) or intrathecally (1 × 10⁶ cells per 50 µL). B6.SJL-Ptprca Pepcb/BoyJ mice were 6 to 10 weeks of age when spleens were extracted to generate CAR T cells. B6;C3-Tg(Prnp-MAPT*P301S)PS19Vle/J mice were 8 to 10 weeks of age when treated with CAR T cells intrathecally, then aged to 9 months for behavioral testing. Following experimental procedures, mice were either sacrificed for further analyses or placed into a survival cohort and monitored until reaching a humane endpoint. The animals were housed in a temperature- and humidity-controlled environment under a 12-h light-dark cycle with regular rodent chow and sterilized water. All strains included both sexes, with individuals randomized by sex across experimental groups. All experiments were approved by the CSHL Institutional Animal Care and Use Committee.

### Isolation, expansion and transduction of mouse T cells

B6.SJL-Ptprca Pepcb/BoyJ mice were euthanized and spleens were collected. Following tissue dissection and red blood cell lysis using ACK Lysing Buffer (Thermo Fisher, A1049201), primary mouse T cells were purified using the Pan T Cell Isolation Kit II, mouse (Miltenyi Biotec, 130-095-130). Purified T cells were cultured in RPMI 1640 Medium (Fisher Scientific, 11-875-093) supplemented with 10% FBS (Corning, 35-010-CV), 10 mM HEPES (Thermo Fisher, 15630106), 2 mM L-glutamine (Thermo Fisher, 25030081), 1× MEM Non-Essential Amino Acids Solution (Fisher Scientific, 11-140-050), 55 µM 2-mercaptoethanol (Thermo Fisher, 31350010), 1 mM sodium pyruvate (Thermo Fisher, 11360070), 200 IU/mL recombinant mouse IL-2 (PeproTech, 212-12-500UG), and Dynabeads Mouse T-Activator CD3/CD28 (Thermo Fisher, 11452D) at a bead:cell ratio of 1:2. T cells were spinoculated with retroviral supernatant 24 h after initial T cell activation, as previously described^36,37^ and used for functional analysis 3–4 days later.

### Busulfan conditioning and bone marrow transplantation

Recipient female mice were 17–22-month-old B6.SJL-Ptprca Pepcb/BoyJ (CD45.1) mice obtained from The Jackson Laboratory and aged in-house. Sub-lethal myeloablative conditioning was achieved by intraperitoneal administration of busulfan (20 mg/kg/day for 3 consecutive days; Fisher Scientific, AAJ6134806), beginning 7 days prior to transplantation. On the day of transplantation, myeloid-biased HSPCs (see “Hematopoietic lineage depletion and myeloid-biased HSPC culture” below) were resuspended at 1,000 cells/µL in PBS, and 200 µL (2 × 10⁵ cells) was injected per recipient via the tail vein without co-transplantation of carrier/support cells, using a rotating tail-vein injection restrainer.

### Brain isolation and processing

Mice were sacrificed via intraperitoneal injection of 250 mg/kg 2,2,2-tribromoethanol (Sigma-Aldrich, T48402-25G) prior to transcardial perfusion with PBS. Brains were extracted, dural tissue was removed, and the brains were minced with a razor blade and enzymatically digested using the Neural Tissue Dissociation Kit (P) (Miltenyi Biotec, 130-092-628) on a gentleMACS Octo Dissociator. Digestion was quenched with 10 mg/mL ovomucoid trypsin inhibitor (Sigma-Aldrich, T9253-250MG) and 10 mg/mL bovine serum albumin (Fisher Scientific, 50-253-893) prepared in HBSS, no calcium, no magnesium, no phenol red (Fisher Scientific, 14-175-095). The resulting suspension was filtered through a 70 µm filter and washed with 20 mL HBSS. Cells were resuspended in 30% isotonic Percoll (Sigma-Aldrich, P4937-500ML) and centrifuged at 700 × g for 15 min. Red blood cells were lysed using ACK Lysing Buffer (Thermo Fisher, A1049201) for 5 min. Cells were resuspended in FACS buffer (1× HBSS, no calcium, no magnesium, no phenol red [Fisher Scientific, 14-175-095] + 2% FBS [Corning, 35-010-CV]).

### Bone marrow isolation and processing

Lower limbs were harvested from euthanized mice following sterilization of the abdomen and hind legs with 70% ethanol. A midline incision was made, and the skin was reflected to expose the hind limbs, which were removed at the hip joint; the femur, tibia, and hip bones were dissected free of surrounding soft tissue, and feet were removed with the femur and tibia separated at the knee joint. Muscle and connective tissue were carefully trimmed from the bones, which were kept intact throughout. Bone marrow was extracted from cleaned bones using one of two methods. In the first, marrow was flushed from the bone cavity with FACS buffer (1× HBSS, no calcium, no magnesium, no phenol red [Fisher Scientific, 14-175-095] + 2% FBS [Corning, 35-010-CV]) using a syringe and needle, and the resulting suspension was pelleted by centrifugation (8,000 rpm, 1 min). In the second, femoral and tibial epiphyses were removed with fine scissors, and intact bones were placed cut-end-down into a perforated 0.6 mL microcentrifuge tube nested within a 1.5 mL microcentrifuge tube; centrifugation at 10,000 × g for 15 s pelleted the marrow directly into the outer tube. Following extraction by either method, the supernatant was discarded and the marrow pellet was resuspended in ACK Lysing Buffer (Thermo Fisher, A1049201) to lyse red blood cells; the reaction was quenched with FACS buffer after 5 min at room temperature. Cells were filtered through a 40 µm strainer and pelleted by centrifugation at 300 × g for 5 min, the centrifugation condition used for all subsequent steps unless otherwise noted. Cells were resuspended in FACS buffer and counted.

### Hematopoietic lineage depletion for bone marrow derived macrophage (BMDM) differentiation

Hematopoietic lineage depletion was performed on ACK-lysed whole bone marrow using a magnetic bead-based system (Lineage Cell Depletion Kit, mouse; Miltenyi Biotec, 130-110-470) according to the manufacturer’s instructions. Briefly, per 2 × 10⁷ cells, the pellet was resuspended in 80 µL MACS buffer and 20 µL Lineage Cell Depletion Cocktail was added, followed by a 10 min incubation on ice. Columns were equilibrated with 3 mL MACS buffer, and cells were brought to 1 mL total volume and loaded onto the column. The column was washed three times with 3 mL MACS buffer each, allowing each wash to fully elute before adding the next; the combined flow-through (lineage-depleted fraction) was retained and pelleted by centrifugation (300 × g, 5 min). This pellet represented the lineage-depleted population used for BMDM differentiation or bulk RNA sequencing below.

### BMDM differentiation and culture

Lineage-depleted cells were resuspended in BMDM differentiation medium (DMEM, high glucose, GlutaMAX Supplement, HEPES [Gibco, 10564011] supplemented with 10% FBS [Corning, 35-010-CV], 1% Penicillin:Streptomycin Solution [Fisher Scientific, 50-753-3040], and 25 ng/mL recombinant murine M-CSF [PeproTech, 315-02-10UG]; final dilution 1:2000 from a 50 µg/mL stock) and seeded at 2 × 10⁵ cells per well in 2 mL medium in 6-well plates. A half-medium change with fresh M-CSF-supplemented medium was performed on day 3–4. Cells were considered fully differentiated into BMDMs after 7 days of culture. Culture medium was collected, normalized to total RNA concentration, and processed and measured by Eve Technologies using the Mouse Cytokine/Chemokine 68-Plex Discovery Assay® Array (MD68) for cytokine analysis. Cells were collected for downstream analysis.

### Hematopoietic lineage depletion and myeloid-biased HSPC culture

For transplantation experiments, ACK-lysed whole bone marrow was subjected to hematopoietic lineage depletion using the Lineage Cell Depletion Kit, mouse (Miltenyi Biotec, 130-110-470) according to the manufacturer’s instructions, to enrich for hematopoietic stem and progenitor cells (HSPCs). Isolated HSPCs were cultured overnight in StemSpan Serum-Free Expansion Medium (STEMCELL Technologies, 09650) supplemented with 10 ng/mL each of recombinant murine M-CSF (R&D Systems, 414-CS-005), G-CSF (R&D Systems, 415-ML-010), and IL-6 (PeproTech, 216-16-10UG) to induce myeloid lineage bias. The following day, cells were harvested, pooled by experimental condition (3–4 mice per condition), and counted using an automated cell counter. Cell suspensions were adjusted to 1,000 cells/µL in PBS for transplantation.

### Immunofluorescence staining

Mice were sacrificed via intraperitoneal injection of 250 mg/kg 2,2,2-tribromoethanol (Sigma-Aldrich, T48402-25G) prior to transcardial perfusion with PBS. Brains were extracted and cut sagittally, then immersion-fixed in 4% PFA at 4°C for 48 h. Fixed brains were subsequently processed through a graded ethanol series, cleared, and embedded in paraffin blocks. Paraffin-embedded brains were sectioned and mounted onto glass slides. Prior to staining, slides were deparaffinized and rehydrated through sequential immersion in Histo-Clear (National Diagnostics, HS-200-5gal) (2 × 5 min), followed by a descending ethanol series (100%, 95%, 70%, 40%) and a final rinse in distilled water. Antigen retrieval was performed by immersing slides in Antigen Unmasking Solution, Citric Acid Based (Vector Laboratories, H-3300-250), and heating in an Instant Pot on high pressure for 15 min, followed by a quick pressure release and cooling to room temperature. Slides were washed 3 × 5 min in PBS-T (0.05% Tween-20; Sigma-Aldrich, P1379-100ML) and blocked in 2% normal donkey serum (NDS; Jackson ImmunoResearch, 017-000-121)/1% bovine serum albumin (BSA; Fisher Scientific, 50-253-893) prepared in PBS-T for 1 h at room temperature. Primary antibodies, diluted in blocking solution, were applied and slides were incubated overnight at 4°C. Slides were washed 3 × 5 min in PBS-T, then incubated with secondary antibodies and DAPI (Sigma-Aldrich, D9542-5MG) diluted in blocking solution for 1 h at room temperature. Slides were washed 3 × 5 min in PBS-T, treated with TrueBlack Plus Lipofuscin Autofluorescence Quencher (Biotium, 23014) for 10 min, and washed 3 × 5 min in PBS (no Tween-20). Slides were mounted using Invitrogen ProLong Gold Antifade Mountant (Fisher Scientific, P36930).

### Immunohistochemical staining

Mice were sacrificed via intraperitoneal injection of 250 mg/kg 2,2,2-tribromoethanol (Sigma-Aldrich, T48402-25G) prior to transcardial perfusion with PBS. Brains were extracted and cut sagittally, then immersion-fixed in 4% PFA at 4°C for 48 h. Fixed brains were subsequently processed through a graded ethanol series, cleared, and embedded in paraffin blocks. Paraffin-embedded brains were sectioned and mounted onto glass slides. Prior to staining, slides were deparaffinized and rehydrated through sequential immersion in Histo-Clear (National Diagnostics, HS-200-5gal) (2 × 5 min), followed by a descending ethanol series (100%, 95%, 70%, 40%) and a final rinse in distilled water. Antigen retrieval was performed by immersing slides in Antigen Unmasking Solution, Citric Acid Based (Vector Laboratories, H-3300-250), and heating in an Instant Pot on high pressure for 15 min, followed by a quick pressure release and cooling to room temperature. Slides were washed 3 × 5 min in PBS-T (0.05% Tween-20; Sigma-Aldrich, P1379-100ML) and blocked using M.O.M. Blocking Reagent (Vector Laboratories, MKB-2213-1), prepared with the normal horse serum (NHS) included in the kit, for 1 h at room temperature. Anti-phospho-Tau (Ser202, Thr205) (clone AT8; Thermo Fisher, MN1020), diluted in blocking solution, was applied and slides were incubated overnight at 4°C. Slides were washed 3 × 5 min in PBS-T, then incubated with M.O.M. ImmPRESS HRP Polymer Kit (Vector Laboratories, MP-2400) for 1 min 30 s at room temperature. Antibody binding was visualized using DAB substrate from the M.O.M. ImmPRESS HRP Polymer Kit (Vector Laboratories, MP-2400), and slides were counterstained with Hematoxylin Stain, Harris Modified (Newcomer Supply, 1201C). Slides were dehydrated through an ascending ethanol series and cleared in Histo-Clear prior to mounting with Permount Mounting Medium (Fisher Scientific, SP15-100).

### Flow cytometry analysis

Single-cell suspensions were generated from bone marrow and brain tissue as described in the “Bone marrow isolation and processing” and “Brain isolation and processing” sections. For all panels except the hematopoietic stem cell (HSC) panel, nonspecific antibody binding was blocked by incubating cell suspensions with anti-CD16/CD32 (BD Biosciences, 553142). Samples were centrifuged and resuspended in FACS buffer supplemented with the panel-appropriate viability dye: Ghost Dye UV 450 (Cytek Biosciences, 13-0868-T100) for the CAR T cell and myeloid cell panels; Ghost Dye Red 780 (Cytek Biosciences, 13-0865-T100) or Zombie Yellow Viability Kit (BioLegend, 423103) for the HSC panel; or Ghost Dye Violet 540 (Cytek Biosciences, 13-0879-T100) for the MPP2/MPP3 sorting panel. Samples were stained with the viability dye for 30 min on ice, then washed with FACS buffer. Samples were subsequently stained with panel-appropriate cell surface antibodies for 45 min on ice, then washed twice with FACS buffer. Following surface staining, samples were either fixed using BD Cytofix (BD Biosciences, 554655) or kept on ice for immediate analysis. Fixed samples were analyzed on either an LSRFortessa flow cytometer (BD Biosciences) or an ID7000 Spectral Cell Analyzer (Sony Biotechnology). Live cells were analyzed and sorted using either a BD FACSymphony S6 cell sorter (BD Biosciences) or an SH800S Cell Sorter (Sony Biotechnology). Sorted cells were either lysed directly in NEBNext lysis buffer (New England Biolabs, E6420S) or TRI Reagent (Zymo Research, R2050-1-200), or collected into PBS or BMDM differentiation medium for downstream processing.

### Protein extraction for cytokine analysis

Snap-frozen mouse brains were thoroughly crushed to a fine powder using a mortar and pestle under liquid nitrogen. The crushed tissue was transferred to a 1.5 mL Eppendorf tube and homogenized in lysis buffer (20 mM Tris-HCl [prepared in-house], 0.5% Tween-20 [Sigma-Aldrich, P1379-100ML], 150 mM sodium chloride [Fisher Scientific, S271-3], and protease inhibitor [Thermo Fisher Scientific, 87786] at a 1:100 dilution). Samples were centrifuged at 15,000 rpm for 10 min at 4°C, and the resulting supernatant was transferred to a fresh tube. Protein concentration was quantified using the Pierce BCA Protein Assay Kit (Thermo Fisher, 23227). Samples were normalized to total protein concentration and processed and measured by Eve Technologies.

### Single-cell RNA sequencing

Single-cell suspensions were generated from bone marrow and brain tissue as described in the “Bone marrow isolation and processing” and “Brain isolation and processing” sections. For single-cell RNA sequencing (scRNA-seq) analysis of uPAR+ and uPAR− sorted brain cells, one sample consisted of 6 pooled aged female mice and the other sample consisted of 6 pooled aged male mice. Prior to sorting on an SH800S Cell Sorter (Sony Biotechnology), these samples were stained with DAPI (Sigma-Aldrich, D9542-5MG) and uPAR PE (clone 109801) (R&D Systems, FAB531P). For all other scRNA-seq analyses, samples were composed of the following pooled groups: 2 young UT female mice, 2 young UT male mice, 2 young uPAR female mice, 2 young uPAR male mice, 2 aged UT female mice, 2 aged UT male mice, 2 aged uPAR female mice, and 2 aged uPAR male mice. Samples were processed using either Chromium Next GEM Single Cell 3’ GEM, Library & Gel Bead Kit v3.1 (10x Genomics, PN-1000121) or Chromium GEM-X Single Cell 3’ Kit v4 (10x Genomics, PN-1000691) chemistry, according to the manufacturer’s instructions. Sequencing reads were aligned to the appropriate 10x Genomics mouse reference genome using Cell Ranger (10x Genomics).

Three mouse single-cell RNA-sequencing (scRNA-seq) experiments were analyzed: (1) FACS-sorted uPAR-positive and uPAR-negative cells from aged untreated brain; (2) whole brain from young and old mice after intrathecal UT or m.uPAR-m.28z CAR T-cell treatment; and (3) progenitor-enriched bone marrow from young and old mice after intravenous UT or m.uPAR-m.28z treatment. Public human datasets included the Siletti et al. adult brain atlas^19^ (four donors), the FreshMG scRNA-seq cohort from Lee et al.^20^ (12 donors), the PsychAD snRNA-seq cohort from Lee et al. ^20^(648 donors: 214 controls and 434 individuals with Alzheimer disease), and healthy CD34-positive marrow from Ainciburu et al.^52^ (GSE180298; eight donors: five young and three elderly). Mouse sequencing reads were processed with 10x Genomics Cell Ranger (v9.0.0) to generate filtered feature-barcode matrices, which were analyzed independently in R (v4.5.0) using Seurat (v5.4.0^38^). Quality-control thresholds were selected after inspection of mitochondrial-read, total-count and detected-gene distributions. Cells were retained using the following thresholds, respectively: sorted brain, <15%, 500–20,000 and 250–5,000; post-treatment brain, <5%, 500–20,000 and 250–5,000; and bone marrow, <10%, 500–60,000 and 250–6,000. Mouse samples were normalized with *SCTransform* for integration. *RunPCA*, *FindNeighbors*, *FindClusters* and *RunUMAP* were used for dimensionality reduction, graph-based clustering and visualization. Sorted-brain samples were integrated using *SelectIntegrationFeatures*, *PrepSCTIntegration*, *FindIntegrationAnchors* and *IntegrateData*, whereas post-treatment brain and marrow used reciprocal-PCA integration. Brain cell labels were assigned with Allen Institute MapMyCells^18,39^ using the 10x Whole Mouse Brain taxonomy (CCN20230722) and consolidated into 14 populations; marrow labels were transferred from the Baccin et al. reference atlas^40^. For the public human datasets, published annotations and embeddings were retained without new SCTransform integration or reclustering. ALRA^41^ was used for PLAUR expression plots. Module scoring, normalized-expression averages, PLAUR-positive versus PLAUR-negative classification and differential expression used observed, non-imputed expression. PLAUR-positive cells were defined by detectable observed normalized expression (>0). Human genes were converted to one-to-one mouse orthologs where required. Across all mouse and human datasets, RNA counts used for expression analyses were normalized with Seurat’s NormalizeData function using the LogNormalize method and a scale factor of 10,000. Module scores were calculated before cell-type subsetting using AddModuleScore. To assess inflammatory and age-associated transcriptional programs across datasets, we evaluated four predefined gene signatures: the Hickman et al. myeloid inflammatory signature^42^, the MSigDB Hallmark Inflammatory Response gene set^43^, the Chambers et al. aging-HSC inflammaging signature^44^ and the Kovtonyuk et al. myeloid-bias signature^45^. Differential expression was performed with FindMarkers using two-sided cell-level Wilcoxon rank-sum tests on observed LogNormalized RNA values. Genes detected in at least 10% of either group with nominal P < 0.05 and absolute average log2 fold change >0.5 were separated by direction for enrichment analysis with enrichR (v3.4^43^).

### Bulk RNA sequencing

Total RNA was isolated using the Direct-zol RNA Purification Kit, Microprep (Zymo Research, R2062) for cell lines or the NEBNext Single Cell/Low Input RNA Library Prep Kit for Illumina (New England Biolabs, E6420S) for ultra-low RNA concentrations from sorted MPP2 and MPP3 cells. Sequencing and library preparation were performed at the Cold Spring Harbor Laboratory (CSHL) Sequencing Core. Libraries were prepared from 120 ng of total RNA using the KAPA mRNA HyperPrep Kit (Fisher Scientific, 50-196-5251) or the NEBNext Single Cell/Low Input RNA Library Prep Kit for Illumina (New England Biolabs, E6420S), following the manufacturer’s standard protocol. Samples were sequenced on an Illumina NextSeq 2000 using a P2 flow cell with paired-end 101 bp reads (PE101) for 200 cycles using NextSeq 2000 Control Software v1.5.0. The resulting RNA-seq data were analyzed by removing adaptor sequences using Cutadapt^46^. RNA-seq reads were then aligned with STAR^47^, and transcript counts were quantified using featureCounts^48^ to generate a raw-count matrix. Differential gene expression (DEG) analysis and adjustment for multiple comparisons were performed using DESeq2^49^, which models raw counts using a negative binomial generalized linear model with gene-wise dispersion estimates shrunk toward a fitted mean-dispersion trend, comparing experimental conditions with at least two independent biological replicates per condition. Statistical significance was assessed using the Wald test, implemented in R Studio (Posit Software). For gene set enrichment analysis (GSEA), genes were ranked by the Wald statistic-derived log2 fold change estimates, which incorporate both the magnitude and precision of the estimated effect size. For over-representation analysis (Enrichr), log2 fold changes were shrunk using the adaptive shrinkage method (ashr^50^) via DESeq2’s lfcShrink() function; associated p-values and adjusted p-values were retained from the original Wald test. Preranked GSEA was performed using clusterProfiler^51^ against Gene Ontology Biological Process and Molecular Signatures Database (MSigDB) Hallmark gene sets, using mouse-native or ortholog-mapped gene sets as appropriate (msigdbr). Over-representation analysis of upregulated and downregulated gene sets was performed using Enrichr against the same gene set collections. For lollipop visualization of GSEA results, pathways were plotted with the normalized enrichment score (NES) on the x-axis, gene set size represented by point size, and statistical significance (−log10 p-value) represented by point color. For lollipop visualization of over-representation analysis results, pathways were plotted with the log odds ratio on the x-axis, combined score represented by point size, and statistical significance (p-value) represented by point color. All graphs were generated in R Studio using the ggplot2 package.

### Morris water maze (MWM)

Spatial learning and reference memory were assessed using the Morris water maze, adapted from^52^. The apparatus consisted of a circular pool filled with water maintained at 23 ± 2°C and rendered opaque with non-toxic white tempera paint. High-contrast spatial cues were affixed to the surrounding walls and remained fixed throughout testing. The pool was divided into four virtual quadrants (NW, NE, SW, SE), with a submerged circular escape platform (10 cm diameter, 1.5 cm below the water surface) positioned in the target quadrant. One day before acquisition training, mice were habituated to swimming and platform escape via three trials in clear water with the platform placed centrally. During the 4-day acquisition phase, mice received four training trials per day, entering the pool facing the wall from one of four starting locations equidistant from the platform quadrant; mice were given a maximum of 60 s to locate the platform and were gently guided to it if unsuccessful, then allowed to remain on the platform for 15 s before being dried and returned to a heated holding cage between trials. Latency to platform, swim speed, path length, and percent time in the target quadrant were recorded using automated video tracking (Noldus, EthoVision XT 17.5).

### Y-maze spatial recognition memory

Short-term spatial recognition memory was assessed using a two-trial Y-maze paradigm^53^. The apparatus consisted of a Y-shaped maze with three identical enclosed arms (35 cm length × 5 cm width × 10 cm height) diverging at 120°, constructed of opaque, white acrylic. In the acquisition trial (5 min), one arm (“novel”) was blocked with an opaque guillotine door, and mice were placed in the start arm and allowed to explore only the start and “familiar” arms. Mice were then returned to their home cage for a 1 h inter-trial interval. In the retrieval trial (5 min), the block was removed and mice were reintroduced to the start arm with free access to all three arms. The apparatus was cleaned with 70% ethanol between all trials and phases. Time spent in and entries into each arm were recorded using automated video tracking (Noldus, EthoVision XT 17.5), and memory retention was quantified as time spent in the novel arm.

### Novel object recognition (NOR)

Non-spatial declarative memory was assessed using the novel object recognition task^54^. Testing was conducted in a square open-field arena (40 × 40 × 40 cm) over a single day to assess short-term memory. During habituation, mice explored the empty arena freely for 5 min, then were removed and allowed to rest for 10 min. During familiarization, two identical objects (A and A’) were placed symmetrically 10 cm from the walls, and mice were placed in the center of the arena facing away from the objects and allowed to explore for 10 min; exploration was defined as orienting the nose toward an object within ≤2 cm while actively sniffing or touching it, excluding rearing on the object. Mice were then removed and allowed to rest for an additional 10 min before testing. During testing, one familiar object was replaced with a novel object (B) of comparable volume but distinct shape, color, and texture, and mice were allowed 10 min to explore both objects; the left/right position of the novel object was counterbalanced across cohorts. The arena and all objects were cleaned with 70% ethanol between every phase and every mouse. Exploration was recorded using automated video tracking (Noldus, EthoVision XT 17.5), and the discrimination index (DI) was calculated as percentage of (time exploring novel object − time exploring familiar object) / total object exploration time.

### Quantification, reproducibility and statistical analysis

Unless specified, statistical analysis was performed using GraphPad Prism 11 (GraphPad). Data distribution was assumed to be normal, but this was not formally tested. Flow cytometry data were analyzed with FlowJo v10.10.1 (BD Biosciences). Images were analyzed with ImageJ2 (National Institutes of Health) or QuPath^55^. Behavioral video tracking was performed using EthoVision XT 17.5 (Noldus). No statistical methods were used to predetermine sample size in the mouse studies, and no method of randomization was used to assign mice to treatment groups, but groups were balanced by sex except in the bone marrow transplantation experiments. Experiments were repeated in replicates and/or from different subjects in independent experiments. Information on experimental repetition and replicates is provided in the figure legends. Mouse conditions were observed by an operator who was blinded to the treatment groups, in addition to the main investigator, who was not blind to group allocation. Data collection and analysis were not performed blind to the conditions of the experiments. Figures were prepared using BioRender.com for scientific illustrations and Illustrator CC 2022 (Adobe).

---

**Supplementary Figure 1. Safety and profile of intrathecal uPAR CAR T cells.**

(A) Absolute counts of CD45.1 and CD3 double positive cells in the brain that are either CD4^+^ or CD8^+^ as determined by flow cytometry 20 days after intrathecal cell infusion. (n=5 mice per group).

(B) Absolute counts of CD45.1 and CD3 double positive cells in the brain that express CD62L and/or CD44 as determined by flow cytometry 20 days after intrathecal cell infusion. (n=5 mice per group).

(C) Absolute counts of CD45.1 and CD3 double positive cells in the brain that are CD25^+^ as determined by flow cytometry 20 days after intrathecal cell infusion. (n=5 mice per group).

(D) Fold change in body weight 24h before and at various times after cell infusion (IV: n=10 mice for young groups, n=9 mice for old groups; IT: n=10 mice per group).

(E) Fold change in body rectal temperature 24h before and at various times after intravenous cell infusion (IV: n=10 mice for young groups, n=9 mice for old groups; IT: n=10 mice per group).

(F) Absolute counts of CD45.1 and CD3 double positive cells in the brain that are either CD4^+^ or CD8^+^ as determined by flow cytometry 16 months after intrathecal cell infusion. (n=12 mice per group).

(G) Absolute counts of CD45.1 and CD3 double positive cells in the brain that express CD62L and/or CD44 as determined by flow cytometry 16 months after intrathecal cell infusion. (n=12 mice per group).

(H) Exploratory behavior heatmap for novel object recognition task 2.5m after intrathecal cell infusion. (UT young, m.uPAR-m.28z young and UT old: n=12 mice per group; m.uPAR-m.28z old: n=11 mice).

(I) Exploratory behavior heatmap for Y-maze spatial memory task 2.5m after intrathecal cell infusion (UT young, m.uPAR-m.28z young and UT old: n=12 mice per group; m.uPAR-m.28z old: n=11 mice).

(J) Exploratory behavior heatmap for day 4 of Morris water maze test 2.5m after intrathecal cell infusion. (n=6 mice per group).

(K) Exploratory behavior heatmap for novel object recognition task 6m after intrathecal cell infusion. (UT young, m.uPAR-m.28z young, UT old: n=12 mice per group, m.uPAR-m.28z old: n=9 mice).

(L) Exploratory behavior heatmap for Y-maze spatial memory task 6m after intrathecal cell infusion. (UT young, m.uPAR-m.28z young, UT old: n=12 mice per group, m.uPAR-m.28z old: n=10 mice).

(M) Exploratory behavior heatmap for day 4 of Morris water maze test 6m after intrathecal cell infusion. (n=6 mice per group).

(N) Exploratory behavior heatmap for novel object recognition task 16m after intrathecal cell infusion. (UT n=12 mice, m.uPAR-m.28z n=10 mice).

(O) Exploratory behavior heatmap for Y-maze spatial memory task 16m after intrathecal cell infusion. (UT n=11 mice, m.uPAR-m.28z n=10 mice).

(P) Exploratory behavior heatmap for day 4 of Morris water maze test 16m after intrathecal cell infusion. (UT n=11 mice, m.uPAR-m.28z n=10 mice).

(Q) Exploratory behavior heatmap for novel object recognition task 6m after intrathecal cell infusion in PS19 mice. (UT n=6 mice, m.uPAR-m.28z n=8 mice).

(R) Exploratory behavior heatmap for day 4 of Morris water maze test 6m after intrathecal cell infusion in PS19 mice. (UT n=6 mice, m.uPAR-m.28z n=8 mice).

Data are the mean ± s.e.m. (A)-(G). Statistical analysis was performed using Mann-Whitney test (A)-(C) and (F)-(G) or two-way ANOVA (D). Data (A-E) and (H-R) represent one independent experiment or (F-G) two independent experiments.

**Supplementary Figure 2. Identity of uPAR^+^ cells in murine and human brains.**

(A) Surface uPAR expression as determined by flow cytometry on the brains of young (3 months) and old (20 months) mice (n=3 mice per group).

(B) Percentage of uPAR^+^ cells in the brains of 18m old mice that are CD45^+^ or PGP9.5^+^ and representative immunofluorescence (n= 8 mice per group).

(C-D) uPAR^+^ and uPAR^−^ cells from the brain of old (18 months old) mice were FACS sorted and subjected to scRNA-seq (n=12 mice pooled into two replicates).

(C) UMAP of the 29,118 retained aged-brain cells colored by 14 categorical cell identities.

(D) Fraction of cells for each of the different cell types shown in C in uPAR^+^ and uPAR^−^ cells.

(E) Number of CD206^+^ uPAR^+^ cells in the brains of 3 or 18m old mice and representative immunofluorescence (Young n=7, Old n=8).

(F) Number of uPAR^+^ BAMs in the brain and representative immunofluorescence staining of CD206 and uPAR 2.5m after intrathecal cell infusion (Young UT: n=7, young m.uPAR-m.28z and old UT: n=8, old m.uPAR-m.28z: n=9 mice).

(G-J) Brains from young (3 months old) and old (18 months old) mice 2.5m after intrathecal cell infusion were subjected to scRNA-seq.

(G) UMAP of retained post-treatment brain cells colored by 14 categorical cell identities.

(H) Fraction of retained cells assigned to each brain cell type in young UT and old UT mice (n=6 mice pooled into two replicates per group).

(I) Fraction of retained cells assigned to each brain cell type in old UT and old m.uPAR-m.28z mice (old UT: n=6 mice pooled into two replicates; old m.uPAR-m.28z: n=4 mice pooled into two replicates).

(J) Enrichr over-representation analysis^43^ of direction-separated genes differentially expressed between young UT and old UT brains. Odds ratio is shown on the x axis, nominal enrichment P value by color and Enrichr combined score by point size (n=6 mice pooled into two replicates per group).

(K) Number of uPAR^+^ BAMs in the brain and representative immunofluorescence staining of CD206 and uPAR 16m after intrathecal cell infusion (n=6 mice per group).

(L) Heatmap depicting the fold change in the protein levels of proinflammatory cytokines and chemokines in the brains 16 months after intrathecal cell infusion (*n* = 6 mice per group).

(M) Representative immunofluorescence staining of CD206 and uPAR 6m after intrathecal cell infusion in the PS19 mice (UT: n=6 mice; m.uPAR-m.28z: n=7 mice).

(N) Representative immunohistochemistry staining of AT8 Tau 6m after intrathecal cell infusion in the PS19 mice (n=7 mice per group).

(O) Heatmap depicting the fold change in the protein levels of proinflammatory cytokines and chemokines in the brains of PS19 mice 6 months after intrathecal cell infusion (UT n = 5 mice, m.uPAR-m.28z: n=6 mice).

(P) Split-violin plot of ALRA^41^-imputed PLAUR expression in FreshMG cells from young (20-29 years) and old (90+ years) brains^20^. Boxplots show the median and interquartile range. P values were calculated using two-sided cell-level Wilcoxon rank-sum tests (young: n=3,972 cells from one donor; old: n=38,903 cells from 11 donors).

(Q) Enrichr over-representation analysis^43^ of direction-separated genes differentially expressed between observed PLAUR+ and PLAUR-FreshMG cells^20^. Odds ratio is shown on the x axis, nominal enrichment P value by color and Enrichr combined score by point size.

(R) Split-violin plot of ALRA^41^-imputed PLAUR expression in age-matched PsychAD control and Alzheimer disease brains^20^. Boxplots show the median and interquartile range. P values were calculated using two-sided cell-level Wilcoxon rank-sum tests (control: n=35,175 cells from 214 donors; Alzheimer disease: n=85,697 cells from 434 donors).

(S) Enrichr over-representation analysis^43^ of direction-separated genes differentially expressed between observed PLAUR+ and PLAUR-PsychAD cells^20^. Odds ratio is shown on the x axis, nominal enrichment P value by color and Enrichr combined score by point size.

Data are the mean ± s.e.m. (A-B), (F), (K). Statistical analysis was performed using two-tailed unpaired Student’s t-test (A-B), (E), (K); Mann Whitney test (F); two-sided cell-level Wilcoxon rank-sum tests for the ALRA-expression panels (P),(R); and Enrichr over-representation analysis^51^ for panels J, Q and S. For the scRNA-seq enrichment panels, genes detected in at least 10% of either group with nominal P<0.05 and absolute average log2 fold change >0.5 were separated by direction for analysis. Data (A), (E), (K) and represent one independent experiment, or (B), (F) two independent experiments or (M-N) three independent experiments.

**Supplementary Figure 3. Effect of intravenous CAR T cells on uPAR^+^ BAMs.**

(A) Discrimination Index in the novel object recognition task 2.5m after intravenous cell infusion. (UT young, m.uPAR-m.28z young: n=10 mice per group, UT old: n=8 mice; m.uPAR-m.28z old: n=10 mice).

(B) Time spent in novel arm in the Y-maze spatial memory task 2.5m after intravenous cell infusion. (UT young, m.uPAR-m.28z young: n=10 mice per group, UT old: n=8 mice; m.uPAR-m.28z old: n=10 mice).

(C) Latency to the platform in the Morris water maze test 2.5m after intravenous cell infusion. (n=6 mice per group).

(D) Exploratory behavior heatmap for novel object recognition task 6m after intravenous cell infusion. (UT young, m.uPAR-m.28z young: n=10 mice per group, UT old: n=8 mice; m.uPAR-m.28z old: n=10 mice).

(E) Exploratory behavior heatmap for Y-maze spatial memory task 6m after intravenous cell infusion. (UT young, m.uPAR-m.28z young: n=10 mice per group, UT old: n=8 mice; m.uPAR-m.28z old: n=10 mice).

(F) Exploratory behavior heatmap for day 4 of Morris water maze test 6m after intravenous cell infusion. (n=6 mice per group).

(G) Number of uPAR^+^ BAMs in the brain and representative immunofluorescence staining of CD206 and uPAR 6m after intravenous cell infusion (Young UT and young m.uPAR-m.28z: n=8, old UT and old m.uPAR-m.28z: n=14 mice).

(H) Absolute counts of CD45.1 and CD3 double positive cells in the bone marrow that are either CD4^+^ or CD8^+^ as determined by flow cytometry 6 months after intravenous cell infusion. (UT young, m.uPAR-m.28z: n=10 mice, UT old: n=16 mice, m.uPAR-m.28z: n=15 mice).

(I) Absolute counts of CD45.1 and CD3 double positive cells in the bone marrow that express CD62L and/or CD44 as determined by flow cytometry 6 months after intravenous cell infusion. (UT young, m.uPAR-m.28z: n=10 mice, UT old: n=16 mice, m.uPAR-m.28z: n=15 mice).

(J) Absolute counts of monocytes (CD45+ CD3- CD19- NK1.1- CD11b+ Ly6C+ Ly6G-) in the bone marrow as determined by flow cytometry 6 months after intravenous cell infusion. (UT young, m.uPAR-m.28z young: n=10 mice; UT old and m.uPAR-m.28z old: n=16 mice).

(K) uMAP visualization of bone marrow cell types generated by 10X chromium protocol. Colors indicate the 22 different lineages.

(L) Fraction of cells for each of the different cell types for young UT and old UT treated mice (n= 4 mice pooled into two replicates per group).

(M) Fraction of cells for each of the different cell types for old UT and old m.uPAR-m.28z treated mice (n= 4 mice pooled into two replicates per group).

(N) Enrichr over-representation analysis^43^ of direction-separated genes differentially expressed between old UT and young UT bone marrow 6 months after intravenous infusion. Odds ratio is shown on the x axis, nominal enrichment P value by color and Enrichr combined score by point size (n=4 mice per group pooled into two replicates).

Data are the mean ± s.e.m. (A-C), (G-J). Statistical analysis was performed using two-tailed unpaired Student’s t-test (A-C) and (G), Mann-Whitney (H-I), Kruskal-Wallis (J), and Enrichr over-representation analysis^51^ (N). For panel N, genes detected in at least 10% of either group with nominal P<0.05 and absolute average log2 fold change >0.5 were separated by direction for analysis. Data (A-F) and (K-N) represent one independent experiment or (G-J) two independent experiments.

**Supplementary Figure 4. Effects of transplanting uPAR CAR T treated bone marrow progenitors.**

(A) Absolute counts of total CD45.2 CD11b positive cells in the bone marrow as determined by flow cytometry 5 months after bone marrow transplant (UT old to old: n=6 mice, Young to old and old m.uPAR-m.28z to old: n=5 mice).

(B) Absolute counts of total uPAR^+^ CD45.2 CD11b positive cells in the bone marrow as determined by flow cytometry 5 months after bone marrow transplant (UT old to old: n=6 mice, Young to old and old m.uPAR-m.28z to old: n=5 mice).

(C) Absolute counts of total CD45.2^+^ monocytes in the bone marrow as determined by flow cytometry 5 months after bone marrow transplant (UT old to old: n=6 mice, Young to old and old m.uPAR-m.28z to old: n=5 mice).

(D) Absolute counts of total CD45.2^+^ uPAR^+^ monocytes in the bone marrow as determined by flow cytometry 5 months after bone marrow transplant (UT old to old: n=6 mice, Young to old and old m.uPAR-m.28z to old: n=5 mice).

(E) Absolute counts of MHC-II^high^ CD45.2^+^ BAMs in the brain as determined by flow cytometry 5 months after bone marrow transplant (UT old to old: n=6 mice, Young to old and old m.uPAR-m.28z to old: n=5 mice).

(F) Number of uPAR^+^ BAMs in the brain and representative immunofluorescence staining of CD206 and uPAR 5m after bone marrow transplant (UT old to old: n=6 mice, Young to old and old m.uPAR-m.28z to old: n=5 mice).

(G) Absolute counts of total CD45.1 CD11b positive cells in the bone marrow as determined by flow cytometry 5 months after bone marrow transplant (UT old to old: n=6 mice, Young to old and old m.uPAR-m.28z to old: n=5 mice).

(H) Absolute counts of total uPAR^+^ CD45.1 CD11b positive cells in the bone marrow as determined by flow cytometry 5 months after bone marrow transplant (UT old to old: n=6 mice, Young to old and old m.uPAR-m.28z to old: n=5 mice).

(I) Absolute counts of total CD45.1^+^ monocytes in the bone marrow as determined by flow cytometry 5 months after bone marrow transplant (UT old to old: n=6 mice, Young to old and old m.uPAR-m.28z to old: n=5 mice).

(J) Absolute counts of total CD45.1^+^ uPAR^+^ monocytes in the bone marrow as determined by flow cytometry 5 months after bone marrow transplant (UT old to old: n=6 mice, Young to old and old m.uPAR-m.28z to old: n=5 mice).

(K) Absolute counts of total CD45.1^+^ CD11b positive cells in the brain as determined by flow cytometry 5 months after bone marrow transplant (UT old to old: n=6 mice, Young to old and old m.uPAR-m.28z to old: n=5 mice).

(L) Absolute counts of total uPAR^+^ CD45.1 CD11b positive cells in the brain as determined by flow cytometry 5 months after bone marrow transplant (UT old to old: n=6 mice, Young to old and old m.uPAR-m.28z to old: n=5 mice).

(M) Absolute counts of MHC-II^high^ CD45.1^+^ BAMs in the brain as determined by flow cytometry 5 months after bone marrow transplant (UT old to old: n=6 mice, Young to old and old m.uPAR-m.28z to old: n=5 mice).

(N) Absolute counts of uPAR^+^ MHC-II^high^ CD45.1^+^ BAMs in the brain as determined by flow cytometry 5 months after bone marrow transplant (UT old to old: n=6 mice, Young to old and old m.uPAR-m.28 zto old: n=5 mice).

(O) Exploratory behavior heatmap for novel object recognition task 3m after bone marrow transplant. (Young to old and old to old and old m.uPAR-m.28z to old: n=5, old UT to old: n=6 mice).

(P) Exploratory behavior heatmap for Y-maze spatial memory task 3m after bone marrow transplant. (Young to old and old to old and old m.uPAR-m.28z to old: n=5, old UT to old: n=6 mice).

(Q) Split-violin plot of ALRA^41^-imputed PLAUR expression in healthy human bone-marrow HSCs from Ainciburu et al.^56^. Boxplots show the median and interquartile range. P values were calculated using two-sided cell-level Wilcoxon rank-sum tests (young: n=2,876 cells from five donors; elderly: n=16,689 cells from three donors).

(R) Split-violin plot of the MSigDB Hallmark Inflammatory Response gene set^57^ scored with Seurat’s *AddModuleScore* in observed PLAUR^−^ and PLAUR^+^ HSCs from elderly donors in Ainciburu et al. ^56^. Boxplots show the median and interquartile range. P values were calculated using two-sided cell-level Wilcoxon rank-sum tests (PLAUR^−^: n=15,536 cells; PLAUR^+^: n=1,153 cells).

(S) Split-violin plot of ALRA^41^-imputed PLAUR expression in healthy human bone-marrow GMPs from Ainciburu et al. ^56^. Boxplots show the median and interquartile range. P values were calculated using two-sided cell-level Wilcoxon rank-sum tests (young: n=3,174 cells from five donors; elderly: n=926 cells from three donors).

(T) Split-violin plot of the MSigDB Hallmark Inflammatory Response gene set^57^ scored with Seurat’s *AddModuleScore* in observed PLAUR^−^ and PLAUR^+^ GMPs from elderly donors in Ainciburu et al. ^56^. Boxplots show the median and interquartile range. P values were calculated using two-sided cell-level Wilcoxon rank-sum tests (PLAUR^−^: n=842 cells; PLAUR^+^: n=84 cells). Data are the mean ± s.e.m. (A-N). Statistical analysis was performed using two-tailed unpaired Student’s t-test (A-E), (G-N), Mann-Whitney (F), or two-sided cell-level Wilcoxon rank-sum tests with Benjamini-Hochberg correction across related comparisons (Q-T). Data (A-T) represent one independent experiment.

## References

1 2025 Alzheimer’s disease facts and figures. Alzheimer’s & Dementia 21 10.1002/alz.70235

2 van Dyck, C. H. et al. Lecanemab in Early Alzheimer’s Disease. N Engl J Med 388, 9–21 (2023). 10.1056/NEJMoa2212948

3 Sims, J. R. et al. Donanemab in Early Symptomatic Alzheimer Disease: The TRAILBLAZER-ALZ 2 Randomized Clinical Trial. JAMA 330, 512–527 (2023). 10.1001/jama.2023.13239

4 Heneka, M. T. et al. Neuroinflammation in Alzheimer disease. Nat Rev Immunol 25, 321–352 (2025). 10.1038/s41577-024-01104-7

5 Cummings, J. L. et al. Alzheimer’s disease drug development pipeline: 2025. Alzheimers Dement (N Y) 11, e70098 (2025). 10.1002/trc2.70098

6 Vara-Perez, M. & Movahedi, K. Border-associated macrophages as gatekeepers of brain homeostasis and immunity. Immunity 58, 1085–1100 (2025). 10.1016/j.immuni.2025.04.005

7 Da Mesquita, S. & Rua, R. Brain border-associated macrophages: common denominators in infection, aging, and Alzheimer’s disease? Trends Immunol 45, 346–357 (2024). 10.1016/j.it.2024.03.007

8 Mrdjen, D. et al. High-Dimensional Single-Cell Mapping of Central Nervous System Immune Cells Reveals Distinct Myeloid Subsets in Health, Aging, and Disease. Immunity 48, 599 (2018). 10.1016/j.immuni.2018.02.014

9 Bastos, J. et al. Monocytes can efficiently replace all brain macrophages and fetal liver monocytes can generate bona fide SALL1(+) microglia. Immunity 58, 1269–1288 e1212 (2025). 10.1016/j.immuni.2025.04.006

10 Wang, L. et al. CCR2(+) monocytes replenish border-associated macrophages in the diseased mouse brain. Cell Rep 43, 114120 (2024). 10.1016/j.celrep.2024.114120

11 Du, S. et al. Brain-engrafted monocyte-derived macrophages from blood and skull-bone marrow exhibit distinct properties. Neuron 114, 1986–2005 e1989 (2026). 10.1016/j.neuron.2026.01.032

12 Van Hove, H. et al. A single-cell atlas of mouse brain macrophages reveals unique transcriptional identities shaped by ontogeny and tissue environment. Nat Neurosci 22, 1021–1035 (2019). 10.1038/s41593-019-0393-4

13 Amor, C. et al. Senolytic CAR T cells reverse senescence-associated pathologies. Nature 583, 127–132 (2020). 10.1038/s41586-020-2403-9

14 Amor, C., Maestre-Fernandez, I., Chowdhury, S., Ho, Y., Nadella, S., Graham, C., Carrasco, S.E., Nnuji-John, E., Feucht, J., Hinterleitner, C., Barthet, V.J.A., Boyer, J., Mezzadra, R., Wereski, M.G., Tuveson, D.A., Levine, L.R., Jones, L.W., Sadelain, M., Lowe S.W. Prophylactic and long-lasting efficacy of senolytic CAR T cells against age-related metabolic dysfunction. Nat Aging (2024). 10.21203/rs.3.rs-3385749/v1

15 Boskovic, P. et al. Engineering chimeric antigen receptor CD4 T cells for Alzheimer’s disease. Proc Natl Acad Sci U S A 123, e2530977123 (2026). 10.1073/pnas.2530977123

16 Chen, Y. et al. Targeting amyloid-beta pathology by chimeric antigen receptor astrocyte (CAR-A) therapy. Science 391, eads3972 (2026). 10.1126/science.ads3972

17 Yoshiyama, Y. et al. Synapse loss and microglial activation precede tangles in a P301S tauopathy mouse model. Neuron 53, 337–351 (2007). 10.1016/j.neuron.2007.01.010

18 Daniel, S. F. et al. MapMyCells: High-performance mapping of unlabeled cell-by-gene data to reference brain taxonomies. bioRxiv (2026). 10.64898/2026.03.06.710160

19 Siletti, K. et al. Transcriptomic diversity of cell types across the adult human brain. Science 382, eadd7046 (2023). 10.1126/science.add7046

20 Lee, D. et al. Plasticity of Human Microglia and Brain Perivascular Macrophages in Aging and Alzheimer’s Disease. medRxiv (2024). 10.1101/2023.10.25.23297558

21 Sankowski, R. et al. Multiomic spatial landscape of innate immune cells at human central nervous system borders. Nat Med 30, 186–198 (2024). 10.1038/s41591-023-02673-1

22 Rossi, D. J. et al. Cell intrinsic alterations underlie hematopoietic stem cell aging. Proc Natl Acad Sci U S A 102, 9194–9199 (2005). 10.1073/pnas.0503280102

23 Beerman, I. et al. Functionally distinct hematopoietic stem cells modulate hematopoietic lineage potential during aging by a mechanism of clonal expansion. Proc Natl Acad Sci U S A 107, 5465–5470 (2010). 10.1073/pnas.1000834107

24 Hu, M. et al. Senescent-like border-associated macrophages regulate cognitive aging via migrasome-mediated induction of paracrine senescence in microglia. Nat Aging 5, 2039–2054 (2025). 10.1038/s43587-025-00956-5

25 Schonhoff, A. M. et al. Border-associated macrophages mediate the neuroinflammatory response in an alpha-synuclein model of Parkinson disease. Nat Commun 14, 3754 (2023). 10.1038/s41467-023-39060-w

26 Adler, D. et al. Functional border-associated macrophages limit Alzheimer’s Disease progression. bioRxiv (2026). 10.64898/2026.01.31.703045

27 Wu, X., Saito, T., Saido, T. C., Barron, A. M. & Ruedl, C. Microglia and CD206(+) border-associated mouse macrophages maintain their embryonic origin during Alzheimer’s disease. Elife 10 (2021). 10.7554/eLife.71879

28 Cugurra, A. et al. Skull and vertebral bone marrow are myeloid cell reservoirs for the meninges and CNS parenchyma. Science 373 (2021). 10.1126/science.abf7844

29 Abellanas, M. e. a. Alzheimer’s disease causes bone marrow myelopoiesis dysfunction. BioRxiv (2025). 10.1101/2025.10.06.680030

30 Mu, W. C. et al. Trained immunity links hematopoietic stem cell aging to aging-associated inflammation. Nat Aging (2026). 10.1038/s43587-026-01175-2

31 Zeng, A. G. X. et al. Human haematopoietic stem cells remember inflammatory stress. Nature 655, 458–467 (2026). 10.1038/s41586-026-10522-7

32 Tjwa, M. et al. Membrane-anchored uPAR regulates the proliferation, marrow pool size, engraftment, and mobilization of mouse hematopoietic stem/progenitor cells. J Clin Invest 119, 1008–1018 (2009). 10.1172/JCI36010

33 Cetinsoy, O. et al. Gene Association Study of the Urokinase Plasminogen Activator and Its Receptor Gene in Alzheimer’s Disease. J Alzheimers Dis 99, 241–250 (2024). 10.3233/JAD-231383

34 Guo, X. e. a. Causal association of plasminogen activators and their inhibitors with Alzheimer’s disease: a Mendelian randomization study. Arch Med Sci (2024). 10.5114/aoms/192049

35 Holmes, C. Common infections and increased risk of developing dementia: compelling evidence for intervention studies. Lancet Healthy Longev 2, e391–e392 (2021). 10.1016/S2666-7568(21)00147-1

36 Kuhn, N. F. et al. CD40 Ligand-Modified Chimeric Antigen Receptor T Cells Enhance Antitumor Function by Eliciting an Endogenous Antitumor Response. Cancer Cell 35, 473–488 e476 (2019). 10.1016/j.ccell.2019.02.006

37 Davila, M. L., Kloss, C. C., Gunset, G. & Sadelain, M. CD19 CAR-targeted T cells induce long-term remission and B Cell Aplasia in an immunocompetent mouse model of B cell acute lymphoblastic leukemia. PLoS One 8, e61338 (2013). 10.1371/journal.pone.0061338

38 Stuart, T. et al. Comprehensive Integration of Single-Cell Data. Cell 177, 1888–1902 e1821 (2019). 10.1016/j.cell.2019.05.031

39 Yao, Z. et al. A high-resolution transcriptomic and spatial atlas of cell types in the whole mouse brain. Nature 624, 317–332 (2023). 10.1038/s41586-023-06812-z

40 Baccin, C. et al. Combined single-cell and spatial transcriptomics reveal the molecular, cellular and spatial bone marrow niche organization. Nat Cell Biol 22, 38–48 (2020). 10.1038/s41556-019-0439-6

41 Linderman, G. C. et al. Zero-preserving imputation of single-cell RNA-seq data. Nat Commun 13, 192 (2022). 10.1038/s41467-021-27729-z

42 Hickman, S. E. et al. The microglial sensome revealed by direct RNA sequencing. Nat Neurosci 16, 1896–1905 (2013). 10.1038/nn.3554

43 Chen, E. Y. et al. Enrichr: interactive and collaborative HTML5 gene list enrichment analysis tool. BMC Bioinformatics 14, 128 (2013). 10.1186/1471-2105-14-128

44 Chambers, S. M. et al. Aging hematopoietic stem cells decline in function and exhibit epigenetic dysregulation. PLoS Biol 5, e201 (2007). 10.1371/journal.pbio.0050201

45 Kovtonyuk, L. V. et al. IL-1 mediates microbiome-induced inflammaging of hematopoietic stem cells in mice. Blood 139, 44–58 (2022). 10.1182/blood.2021011570

46 Martin, M. Cutadapt removes adapter sequences from high-throughput sequencing reads. EMBnet.jounral 17, 10–12 10.14806/ej.17.1.200

47 Dobin, A. et al. STAR: ultrafast universal RNA-seq aligner. Bioinformatics 29, 15–21 (2013). 10.1093/bioinformatics/bts635

48 Liao, Y., Smyth, G. K. & Shi, W. featureCounts: an efficient general purpose program for assigning sequence reads to genomic features. Bioinformatics 30, 923–930 (2014). 10.1093/bioinformatics/btt656

49 Love, M. I., Huber, W. & Anders, S. Moderated estimation of fold change and dispersion for RNA-seq data with DESeq2. Genome Biol 15, 550 (2014). 10.1186/s13059-014-0550-8

50 Stephens, M. False discovery rates: a new deal. Biostatistics 18, 275–294 (2017). 10.1093/biostatistics/kxw041

51 Yu, G., Wang, L. G., Han, Y. & He, Q. Y. clusterProfiler: an R package for comparing biological themes among gene clusters. OMICS 16, 284–287 (2012). 10.1089/omi.2011.0118

52 Vorhees, C. V. & Williams, M. T. Morris water maze: procedures for assessing spatial and related forms of learning and memory. Nat Protoc 1, 848–858 (2006). 10.1038/nprot.2006.116

53 Dellu, F., Mayo, W., Cherkaoui, J., Le Moal, M. & Simon, H. A two-trial memory task with automated recording: study in young and aged rats. Brain Res 588, 132–139 (1992). 10.1016/0006-8993(92)91352-f

54 Leger, M. et al. Object recognition test in mice. Nat Protoc 8, 2531–2537 (2013). 10.1038/nprot.2013.155

55 Bankhead, P. et al. QuPath: Open source software for digital pathology image analysis. Sci Rep 7, 16878 (2017). 10.1038/s41598-017-17204-5

56 Ainciburu, M. et al. Uncovering perturbations in human hematopoiesis associated with healthy aging and myeloid malignancies at single-cell resolution. Elife 12 (2023). 10.7554/eLife.79363

57 Liberzon, A. et al. The Molecular Signatures Database (MSigDB) hallmark gene set collection. Cell Syst 1, 417–425 (2015). 10.1016/j.cels.2015.12.004

