## Supplementary figures and images for "CAR T cell targeting of inflammatory myeloid progenitors in the bone marrow remodels border-associated macrophages and reverses cognitive aging"

### Supplementary Figure 1

#### SUPPLEMENTARY FIGURE 1

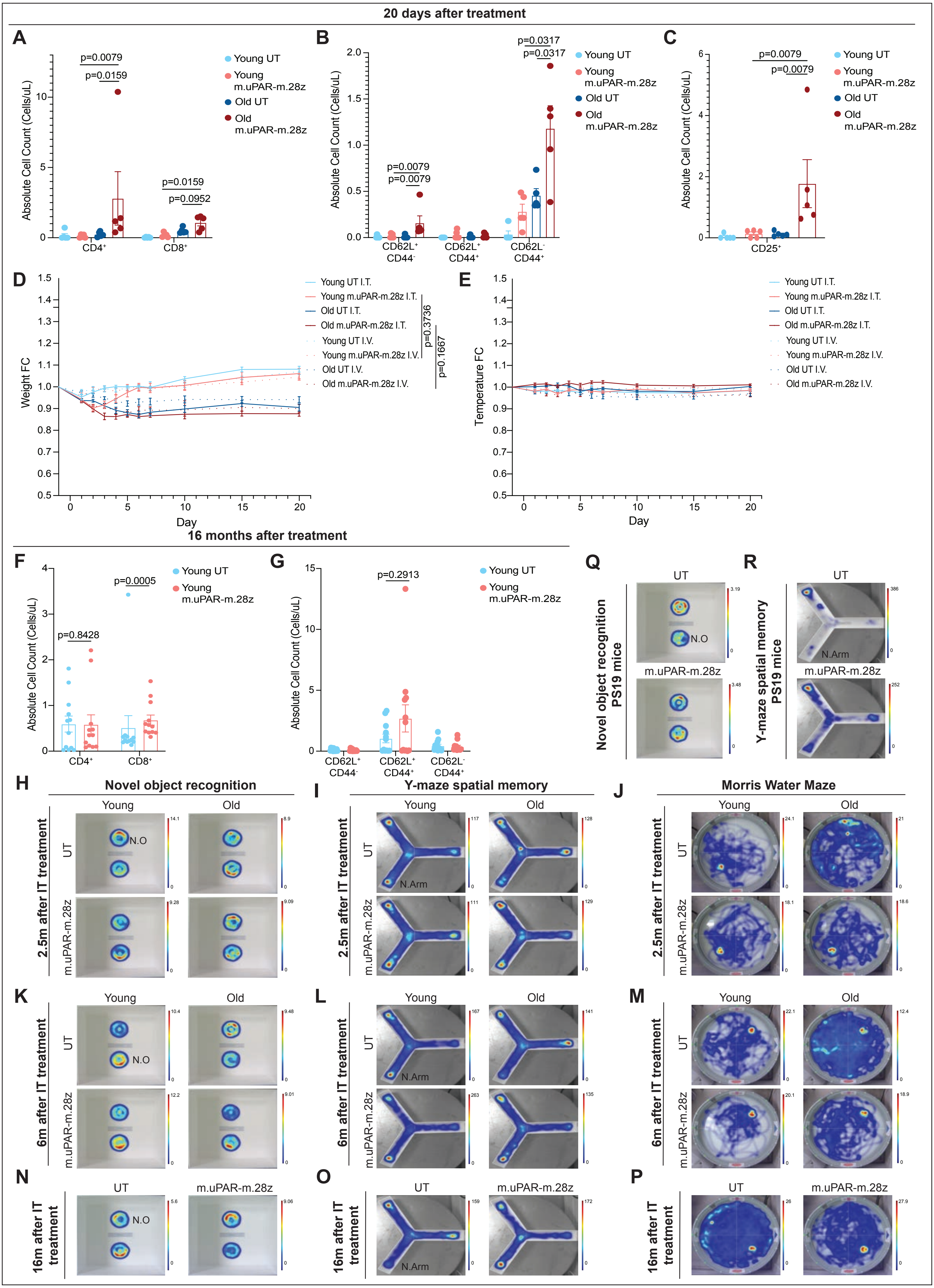

### Supplementary Figure 2

#### SUPPLEMENTARY FIGURE 2

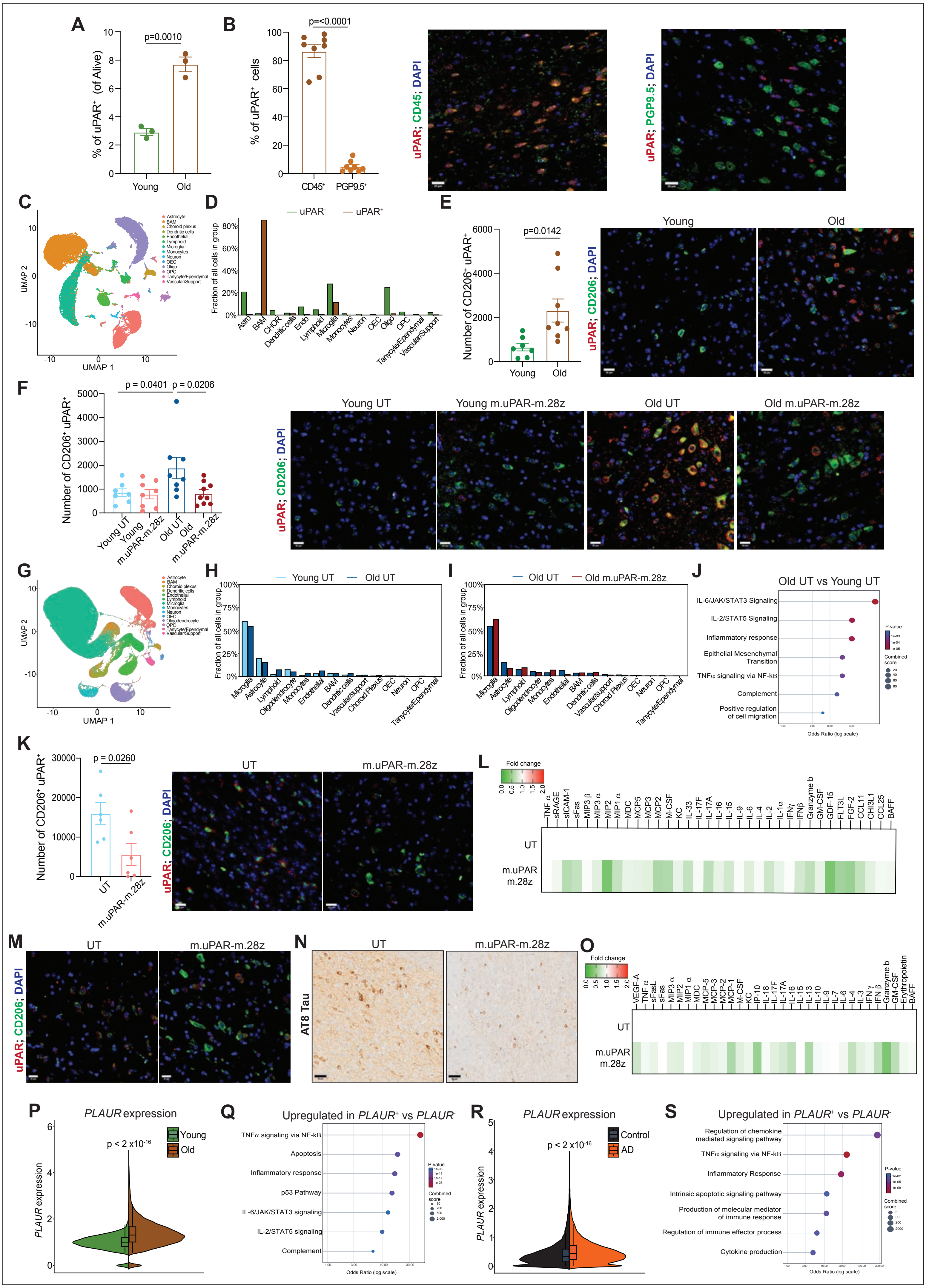

### Supplementary Figure 3

#### SUPPLEMENTARY FIGURE 3

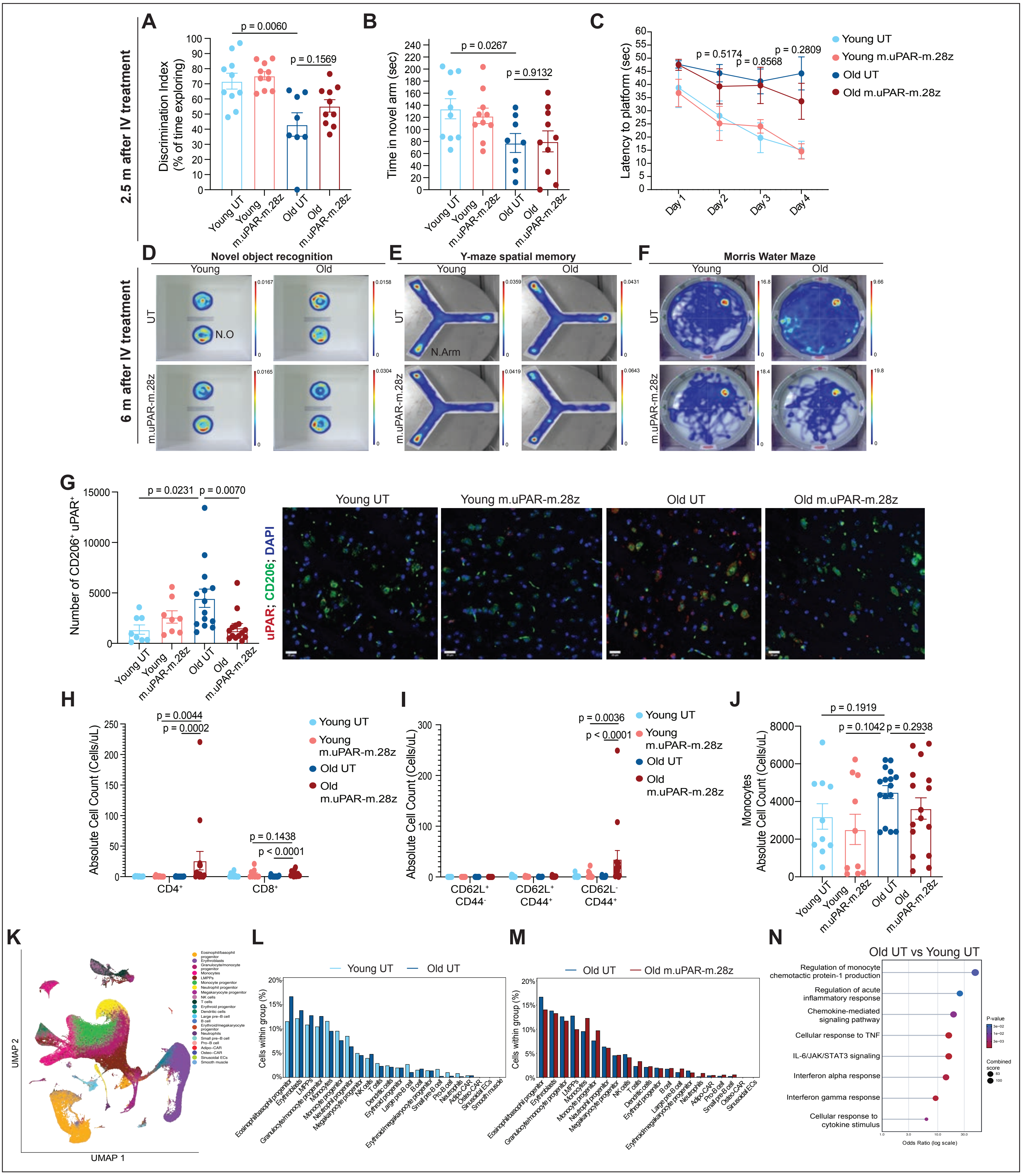

### Supplementary Figure 4

#### SUPPLEMENTARY FIGURE 4

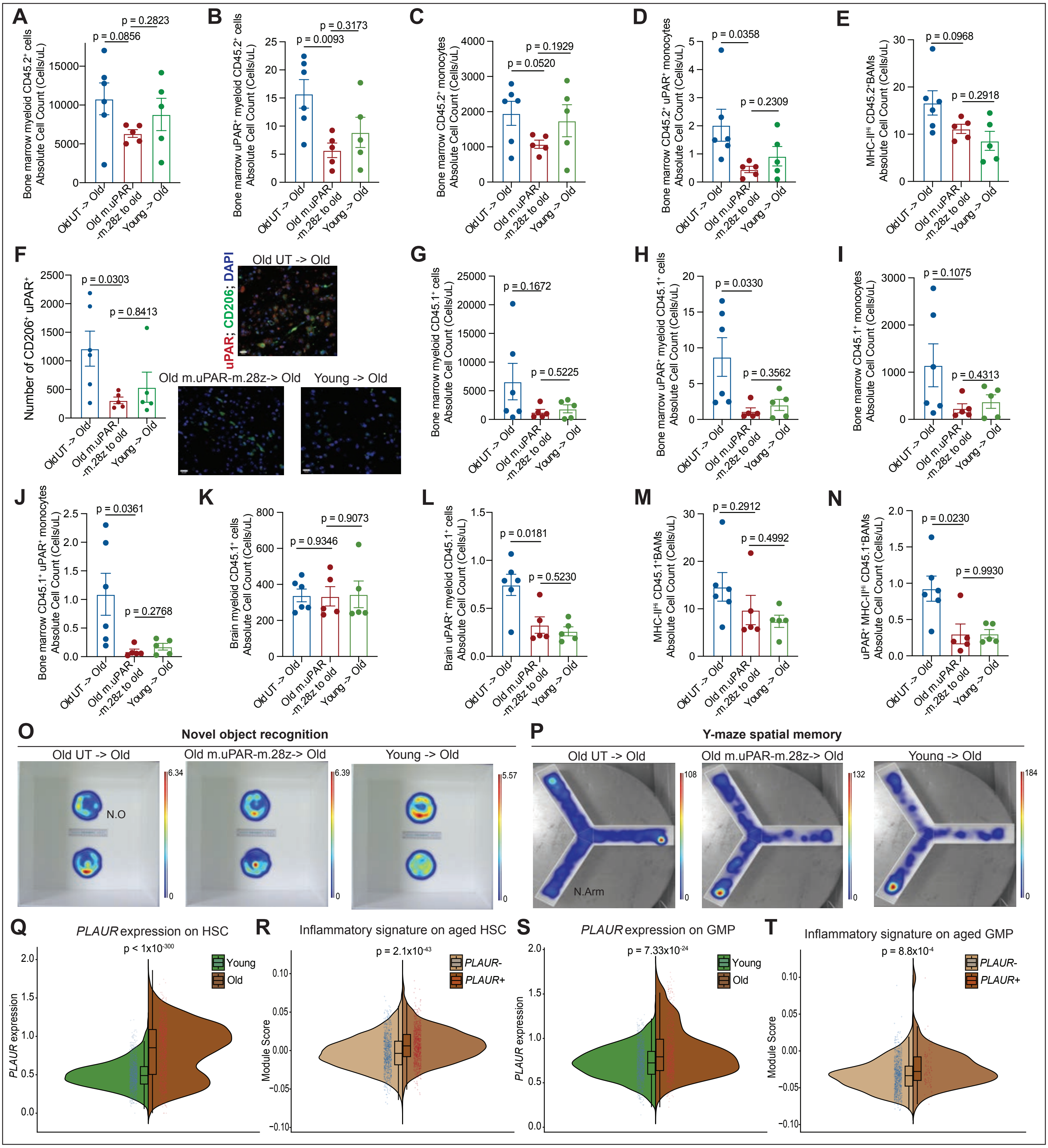
